# Grapevine bacterial communities across the Central Valley of California

**DOI:** 10.1101/2023.07.01.547327

**Authors:** Joel F. Swift, Zoë Migicovsky, Grace E. Trello, Allison J. Miller

**Affiliations:** Department of Biology, Saint Louis University, 3507 Laclede Avenue, St. Louis, MO, 63103, USA; Donald Danforth Plant Science Center, 975 North Warson Road, St. Louis, MO, 63132, USA; Department of Plant, Food and Environmental Sciences, Faculty of Agriculture, Dalhousie University, Truro, NS, Canada; Current address: Kansas Biological Survey & Center for Ecological Research, University of Kansas, Lawrence, KS 66045, USA (J.F.S.); Department of Biology, Acadia University, Wolfville, Nova Scotia, B4P 2R6, Canada (Z.M.)

## Abstract

Plant organs (compartments) host distinct microbiota which shift in response to variation in both development and climate. Grapevines are woody perennial crops that are clonally propagated and cultivated across vast geographic areas, and as such, their microbial communities may also reflect site-specific influences. These site-specific influences, and the microbial differences across site compose ‘terroir’, the environmental influence on wine produced in a given region. Commercial grapevines are typically composed of a genetically distinct root (rootstock) grafted to a shoot system (scion) which adds an additional layer of complexity. In order to understand spatial and temporal patterns of bacterial diversity in grafted grapevines, we used 16S rRNA metabarcoding to quantify soil and compartment microbiota (berries, leaves, and roots) for grafted grapevines in commercial vineyards across three counties in the Central Valley of California over two successive growing seasons. Community composition revealed compartment-specific dynamics. Roots assembled site-specific bacterial communities that reflect rootstock genotype and environment influences, whereas bacterial communities of leaves and berries displayed associations with time. These results provide further evidence of a microbial terroir within the grapevine root systems but also reveal that the microbiota of above-ground compartments are only weakly associated with the local microbiome in the Central Valley of California.

## Introduction

Plants form associations with microorganisms in different compartments across the plant body, including roots, leaves, and fruits. Microbiota vary strongly across compartments due to differences in physical and chemical properties, resource availability, and environmental factors (Martins *et al*. 2013; Hacquard *et al*. 2015; Coleman-Derr *et al*. 2016; Liu *et al*. 2017; Brown *et al*. 2020). Host genetics also play a role in dictating compartment microbiota. For example, root exudate profiles and plant immune responses are often genotype specific and contribute to shaping the plant microbiome (Micallef *et al*. 2009; Mönchgesang *et al*. 2016; Garrido-Oter *et al*. 2018; Sasse *et al*. 2018; Teixeira *et al*. 2019). The associations formed between plants and microorganisms are beneficial to plant survival as microorganisms are able to promote growth and confer resistance to many biotic and abiotic stressors (Xu *et al*. 2018; Durán *et al*. 2018; Rolfe *et al*. 2019). Thus, research has focused on understanding factors that shape a plant’s microbiota, including what stage of development microbial associations form and how stable they are over space and time.

For most plants, the primary source of microorganisms is the surrounding soil (Chi *et al*. 2005; Bulgarelli *et al*. 2013; Zarraonaindia *et al*. 2015; Liu *et al*. 2017). Differences in soil properties, including texture, pH, and chemical composition shape local soil microbiota (Fierer and Jackson 2006; Lauber *et al*. 2009; O’Brien *et al*. 2016; Fierer 2017; Thompson *et al*. 2017). As a result, microbial biogeography is expected to be a primary driver of the plant microbiome (Bokulich *et al*. 2014; Liu *et al*. 2017; Walters *et al*. 2018). We thus hypothesize that the microbiota of each plant compartment reflects, at least in part, the microbiota of the soil in which the plant is growing. Clonally propagated perennial plants offer a unique opportunity to test this hypothesis as the host genotype can be planted across a wide range of soil types to investigate the influence of microbial biogeography on the microbiota of different compartments the plant.

Beyond investigations of the primary source of plant microbiota are questions associated with how the microbiome changes over the life of the host plant. Plant microbiomes are dynamic (Shi *et al*. 2015; Comby *et al*. 2016; Edwards *et al*. 2018; Grady *et al*. 2019; Liu and Howell 2021) and change over development (Micallef *et al*. 2009; Chaparro *et al*. 2014; Torres-Cortés *et al*. 2018). In annual plants, a two-stage model of microbiome change has been proposed. First, seed germination is followed by a period of microbial colonization of the root endosphere and rhizosphere, leading to the establishment of a juvenile microbiome. This juvenile microbiome is composed of an assortment of soil microorganisms in close physical proximity to the seedling as well as endophytes carried by the seed itself (Truyens *et al*. 2015). Second, as the plant reaches reproductive maturity, the juvenile microbiome is displaced by the more stable adult microbiome (Edwards *et al*. 2018). This shift to an adult microbiome is hypothesized to be the result of variation in root exudate composition during a plant’s life cycle (Davey and van Staden 1976; Chaparro *et al*. 2013), which can change rapidly (Zhalnina *et al*. 2018). These data suggest that plant microbiomes reflect changes in plant physiology and exudation profiles over development.

Perennial species may follow the two-stage model of microbiome establishment described above, but the adult microbiome of perennial plants requires additional consideration due to their extended life cycle. Do microbial communities of perennial plants change from one year to the next? Is this change consistent across compartments, or are compartments changing independently across years? In the perennial herb *Arabis alpina*, the abundance of over half of the bacterial taxa of the root endosphere and rhizosphere were significantly altered across 28 weeks of growth, but no significant changes were correlated with developmental stages (flowering vs non-flowering), indicating the microbiota changes with plant age within the first year but is insensitive to developmental stage in this system (Dombrowski *et al*. 2017). A study investigating the microbiome of the perennial mustard *Boechera stricta* across multiple field sites found that the root microbiota shifted over time, with bacterial diversity in the root decreasing over the four years of the experiment, where leaf microbiota were less responsive to the age of the plant and more responsive to the plant genotype (Wagner *et al*. 2016). These data demonstrate compartment-specific patterning between leaves and roots. It remains to be seen whether this pattern in short lived biennial and perennial plants is generalizable to other perennial crop species, which allocate considerably more resources to harvestable organs than wild species (Olsen and Wendel 2013) or to woody perennials, which have substantially longer lifespans (de Witte and Stöcklin 2010; Thomas 2013).

Woody perennial crops offer an excellent system to investigate how microbiota of plant compartments differ among genotypes, across sites, over the course of a growing season, and from one year to the next. Grapevines (*Vitis* spp. L.) are long-lived woody perennials (>20 production years), in which cultivated varieties (cultivars) are clonally propagated and grown across different regions. Studies have found biogeographical patterning to the microbiome of vineyard soils (Burns *et al*. 2015; Gobbi *et al*. 2022) and differences in grapevine microbiota across sites (often termed microbial terroir; Bokulich *et al*. 2014; Zarraonaindia *et al*. 2015; Portillo *et al*. 2016; Mezzasalma *et al*. 2018). However, the stability of these biogeographical patterns in grapevine microbiota across compartments and growing seasons are only beginning to be explored. For instance, in a study of fungal microbiota associated with Pinot Noir in two nearby vineyards, compartments displayed contrasting responses to biogeography and growing season (Liu and Howell 2021). The root compartment fungal diversity strongly varied by site while all compartments (root, leaf, flower, berry) varied across the growing season according to developmental stage (flowering, fruit set, veraison, harvest). These data suggest that the grapevine microbiome is dynamic over time, and that patterns of fungal diversity vary across the vine.

An additional factor affecting the grapevine microbiome is grafting, a common horticultural technique used in viticulture which joins the root and shoot systems of different genotypes together (Mudge *et al*. 2009; Ollat *et al*. 2016). Grafting presents an additional dimension of phenotypic variation, offering an avenue to explore interactions between the soil microbiome, environment, and the genomes of the host and their influence on the microbiota of different compartments of the vine. Recent work has expanded current understanding of interactions between genotypes of rootstock and scion (Cookson and Ollat 2013; Lecourt *et al*. 2015; Migicovsky *et al*. 2019; Gautier *et al*. 2020; Harris *et al*. 2021; Swift *et al*. 2021). While it is clear that grapevine rootstocks exhibit genotype specific rhizosphere effects on root microbiota (D’Amico *et al*. 2018; Marasco *et al*. 2018, 2022; Berlanas *et al*. 2019; Vink *et al*. 2021), less is known about how rootstock genotype influences the microbiota of the leaves and berries of a grafted scion, and how this changes across spatial and temporal scales.

Our work investigated the bacterial communities of grapevine compartments of multiple rootstock/scion combinations across growing seasons, in geographically distinct vineyards, and over multiple years. To do this, we characterized bacterial communities in compartments of rootstock/scion combinations replicated across three commercial vineyards in the Central Valley of California, over the course of two growing seasons. Our objectives were to **1**) assess soil structure, elemental composition, and microbiome across sites, **2**) characterize seasonal and yearly patterns of bacterial communities across vine compartments, and **3)** determine relative contributions of compartment, site, rootstock genotype, and scion genotype to patterns of microorganism community composition across vine compartments.

## Materials and methods

### Experimental Design and Sampling

We sampled vines in three commercial vineyards located along a 177 kilometer north-south transect running through Madera, Merced, and San Joaquin counties in central California (Figure 1A). Each vineyard contained mature grafted vines (>6 years old; Table S1) composed of one of two scions (‘Cabernet Sauvignon’ or ‘Chardonnay’) and one of three rootstocks (‘Freedom’, ‘1103P’, or ‘Teleki 5C’; Figure 1A; Table 1). Each vineyard has a unique vineyard design (Table S1) and unique soil properties (Table S2). Conventional management practices such as mechanical leaf thinning and spray applications of pesticides and fungicides were employed at all vineyards.

**Figure 1.**
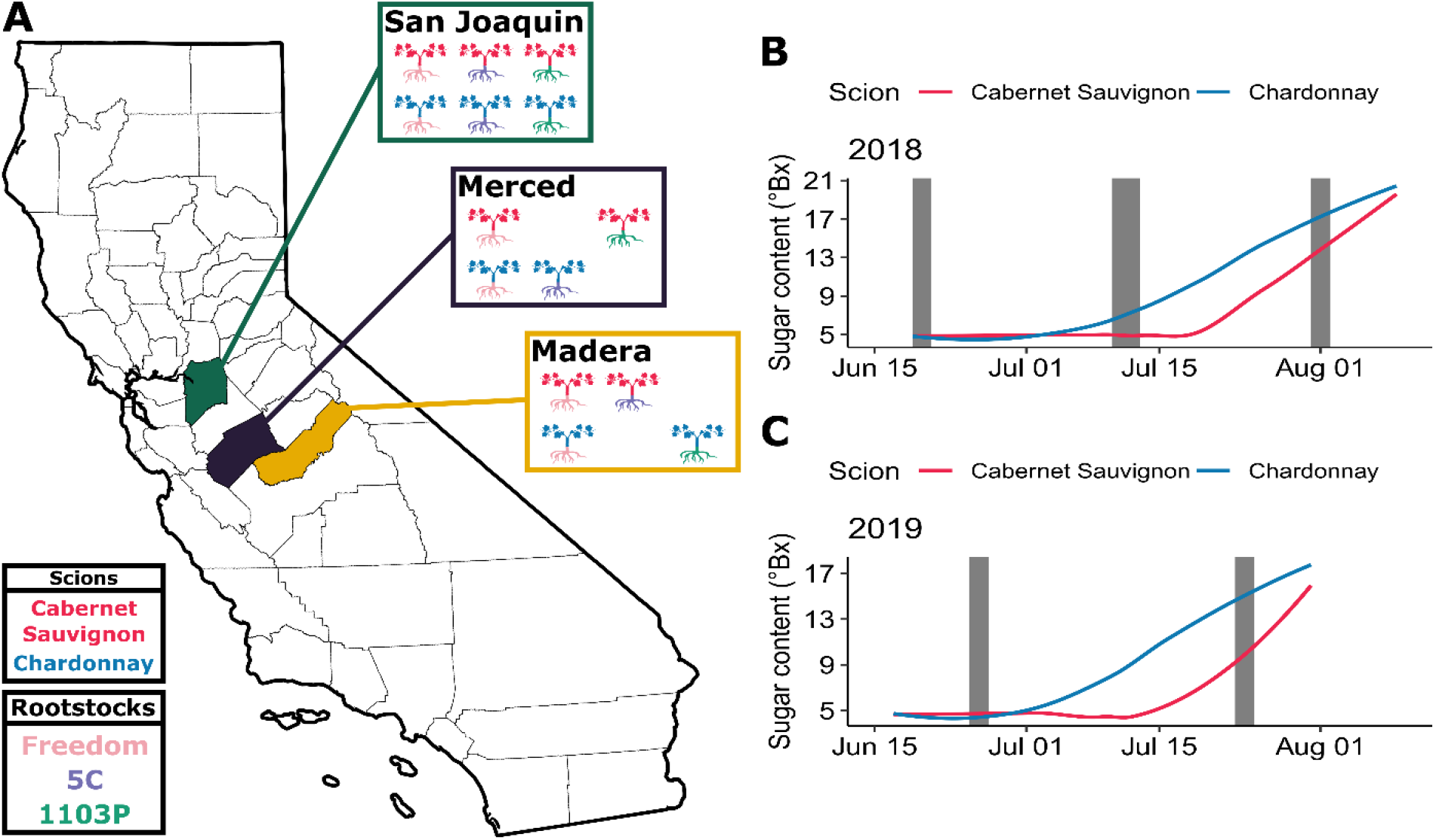
Multiple scion/rootstock combinations were sampled across vineyards and the 2018 and 2019 growing seasons in the central valley of California. **A)** Each of the colored counties represents a vineyard that was sampled. Within each county box (San Joaquin, Merced, and Madera) the scion/rootstock combinations are depicted, corresponding to the legend in the bottom left. The sugar content (measured in degrees Brix) was collected for vines across the **B)** 2018 and **C)** 2019 growing seasons. Grey shading represents collection windows for microbiome sampling.

**Table 1.**
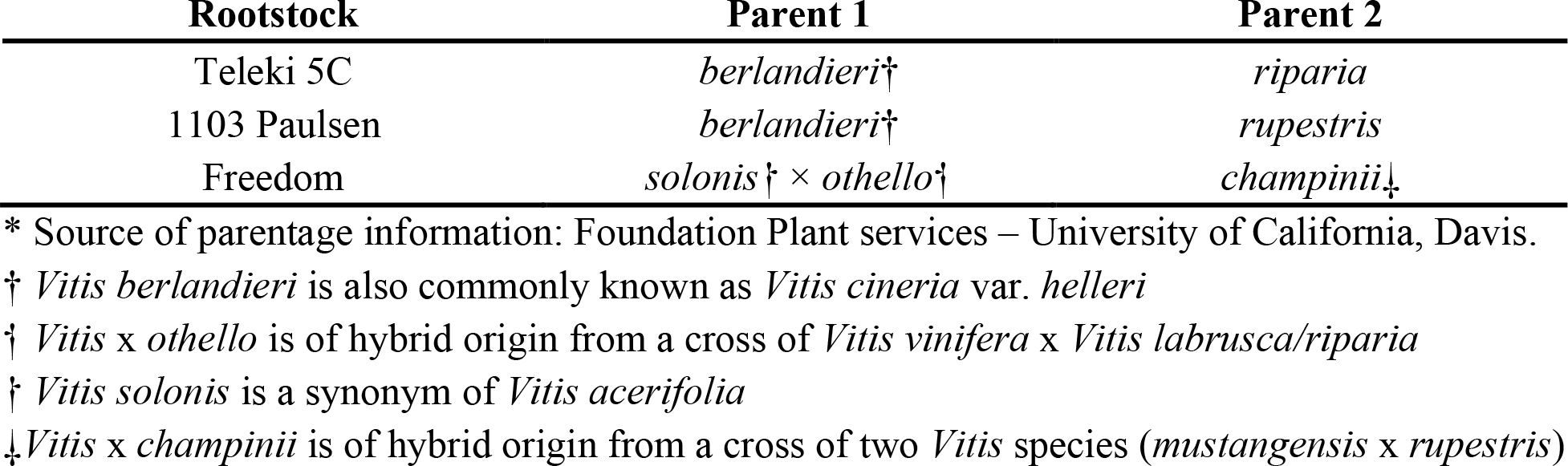
Parentage of rootstocks within this study (Source: Foundation Plant Services - University of California, Davis; http://iv.ucdavis.edu/files/24347.pdf)

One of the three vineyards (San Joaquin) contained all six scion/rootstock combinations (‘Cabernet Sauvignon’ grafted to ‘Freedom’, ‘1103P, and ‘Teleki 5C’; and ‘Chardonnay’ grafted to ‘Freedom’, ‘1103P, and ‘Teleki 5C’). Six collection blocks were sampled at the San Joaquin site (i.e., sets of 24 vines, at least 10 vines from the edge, across 2-3 rows; Figure 1A; Table S1). In the other two vineyards, Madera and Merced, collection blocks for four of the six scion/rootstock combinations were sampled as not all scion/rootstock combinations were present at these vineyards (Figure 1A; Table S1). In Madera, blocks were sampled for ‘Cabernet Sauvignon’ grafted to ‘Teleki 5C’ and ‘Freedom’ and ‘Chardonnay’ grafted to ‘1103P’ and ‘Freedom’. In Merced, blocks for ‘Cabernet Sauvignon’ grafted to ‘1103P’ and ‘Freedom’ and ‘Chardonnay’ grafted to ‘Teleki 5C’ and ‘Freedom’ were established (Figure 1A). From each collection block, three representative vines per scion/rootstock combination were selected. Different vines were sampled in each collection window to minimize disturbance of the vine and its microbiome from repeated collections. Care was taken to ensure that selected vines did not exhibit signs of pathogen infection. We staggered collection windows over the course of the season to capture vine development: in 2018 we made three collections three weeks apart from June 19^th^ – August 2^nd^, 2018, and in 2019 we made two collections four weeks apart June 11^th^ – July 31^st^, 2019 (Figure 1B-C; Figure S1). Sampling periods coincided with multiple developmental stages for the vines, starting at early fruit formation to veraison and, for ‘Chardonnay’, early harvest.

Three compartments (roots, leaves, berries) were sampled from each vine. Roots were collected at a depth of 20-30 cm using a sterile shovel. Three to five leaves approximately 8-12 cm in diameter were collected at roughly the middle position along a shoot and at a height of 1.5 m on the vine. Berries were collected as an intact cluster. We measured total soluble solids (sugar content) in °Brix using a hand-held refractometer (ATAGO) by selecting berries from a damage-free representative cluster on the same vine. Sugar content measurements were collected each time berry clusters were sampled as well as once per week at each vineyard to track berry development (Figure 1B-C). A soil sample was collected during the final sampling time point each year from each collection block. Soil was collected with a sterile shovel, removing the first 3-5cm of topsoil, and passed through a sterile sieve (American Standard No. 16; 1mm pore size). All samples were placed in a cooler with ice packs in the field prior to shipping and storage at -20° C at the Danforth Plant Science Center (St. Louis, MO).

### Soil texture and elemental composition

Soil samples were split into two portions: one for molecular processing (see below) and one for texture and elemental composition analysis. Prior to texture and elemental composition analysis, soil was dried until a consistent weight was achieved, approximately one week at 70°C. For texture analysis, 50g of each soil sample was added to a screw-top jar along with 125mL of sodium hexametaphosphate solution (Sigma-Aldrich; 0.065M) and allowed to agitate overnight on a stir plate. The next day the contents of the jar were transferred to a 1L graduated cylinder and filled to 1L with deionized water. The cylinder was capped and agitated by inverting approximately 30 times for 1 minute. A hydrometer (H-B Instrument Company, 152H) was placed in the cylinder, and the first reading was taken after 40 seconds. Readings were then taken periodically for the next six hours. Elemental composition analysis was conducted at the Agricultural Diagnostic Laboratory (Fayetteville, AR) following established protocols (Sikora and Kissel 2014; Zhang *et al*. 2014) to determine concentrations (in ppm) of the following elements; B, Ca, Cu, Fe, K, Mg, Mn, Na, P, S, and Zn, along with pH.

### DNA extraction and 16S rRNA amplicon sequencing

Soil, root, leaf, and berry samples were processed using previously described methods (Swift *et al*. 2021). DNA extractions were performed with the DNeasy Powersoil Pro Kit (Qiagen) following the manufacturers protocol with two modifications: we used 150mg of plant tissue (non-surface sterilized) per extraction, and we added a 10-minute incubation at 70°C prior to homogenization with a bead mill (Retsch, MM 400). DNA extracts were qualified on a DS-11 Spectrophotometer (DeNovix) and sent to the Environmental Sample Preparation and Sequencing Facility at Argonne National Laboratory for 16S rRNA metabarcoding. Amplicon sequencing and library preparation were conducted following Swift *et al*. (2021). Samples were split into two pools of 336 samples each for sequencing conducted on an Illumina MiSeq, with a 2×151bp Pair-End kit. Samples that produced fewer than 20,000 reads after preliminary quality control and filtering were combined into a third pool that was sequenced on an additional flow cell.

### Amplicon processing and ASV filtering

Sequence processing was conducted using a similar workflow to Swift *et al*. (2021). Briefly, QIIME2 v2021.4 (Bokulich *et al*. 2018) was used to demultiplex samples according to barcode sequence. The DADA2 plugin (Callahan *et al*. 2016) in QIIME2 was used to denoise, dereplicate, and filter chimeric sequences on each sequencing run individually for accurate error model generation. The resulting amplicon sequence variant (ASV) tables and catalogs of representative sequences for each sequence plate were merged. A Naive Bayes classifier pre-trained on the SILVA v.138 16S database (Yilmaz *et al*. 2014; Bokulich *et al*. 2018, 2021) was used for taxonomic classification. ASVs not assigned to a phylum were removed along with ASVs assigned to chloroplasts and mitochondria. The Decontam v1.12.0 package (Davis *et al*. 2017) was used to remove contaminants using the prevalence-based detection method with a threshold of 0.5, removing contaminant ASVs more prevalent in the negative controls than real samples. The data set was filtered to remove singletons by retaining only ASVs present in five or more samples and by removing samples with a read count less than 1,000.

### Statistical analysis

All analyses were conducted within the R environment v4.1.0 (R Core Team 2021). We first modeled the sugar content of the berries of the vines across the growing season. Using a linear model, we assessed effects from the experimental design (rootstock, scion, site, collection week, and their interactions) on sugar content (°Brix). The car package v3.0-11 (Fox and Weisberg 2019) was used to assess the model under a type-3 ANOVA framework.

For soil texture, hydrometer readings were processed using the envalysis package v0.5.1 (Steinmetz 2021) to obtain percentages of sand, silt, and clay. Independent linear models for sand, silt, and clay were fit with collection site and year as main effects. For elemental composition analysis, concentration more than five standard deviations from the mean were removed. A biplot was generated using the factoextra package v1.0.7 (Kassambara and Mundt 2020) to visualize clustering of soil samples by collection site along with the loadings of the principal component analysis (PCA). A linear model was fit to the first two principal components (PC) with collection site and year as main effects. Each of the linear models was assessed via a type-2 ANOVA framework.

ASV counts were normalized by applying a variance stabilizing transformation from the package DESeq2 (Love *et al*. 2014) with a model containing all of main effects (Rootstock + Scion + Compartment + Year + Site + Sugar Content). Alpha diversity statistics were calculated using vegan v2.5.1 (Oksanen *et al*. 2019) and picante v1.8.2 (Kembel *et al*. 2010) in the case of Faith’s Phylogenetic distance (Faith 1992). Linear mixed models were fit via the lmerTest package v3.1.3 (Kuznetsova *et al*. 2017) to assess the effects of the experimental design on alpha diversity indices (response ∼ Rootstock (R) + Scion (S) + Compartment (C) + Sugar Content (Su) + Year + Site + R×S + C×Su). Specific pairwise contrasts from significant terms in the model were further investigated while correcting for multiple comparisons using the emmeans package v.1.6.2.1 (Lenth *et al*. 2020).

We used principal coordinate analysis (PCoA) with Bray-Curtis dissimilarity in order to visualize clustering of samples across experimental factors. PERMANOVA analyses were conducted using the function *Adonis* from the package vegan. For each plant compartment (berries, leaves, roots) a model with Bray-Curtis dissimilarity as the response was fit with all factors as marginal fixed effects, using 1,000 permutations per model. Using a linear model framework, we examined the abundance of each of the top ten bacterial phyla by relative abundance. A linear model was fit with rootstock, scion, brix, year, and site as fixed effects with each phylum as the response variable. The *P-*values from all tests, across phyla, were corrected for multiple testing using false discovery rate (Benjamini and Hochberg 1995).

### Machine learning

For machine learning, categorical factors are preferred over continuous factors to allow for easier statistical interpretation. As such, we discretized berry sugar content values. To choose where to split sugar content values into groups, we plotted values and chose the natural break point in the values. Samples were given the labels pre-ripening (3-7°Bx; n=357) and ripening (>7°Bx; n=237; Figure S2).We used ranger’s v0.13.1 (Wright and Ziegler 2017) implementation of the random forest algorithm in the caret package v6.0.90 (Kuhn 2008) to assess predictability of sample labels and identify ASVs that contribute to predication accuracy. For training the random forest classifier, the dataset was randomly split into a training set (80%) and a testing set (20%). Optimal hyperparameters for each classifier were determined using a grid search over the number of trees (1-501), minimum node size (1, 5, 10), and number of features available at each node (10-100% of the ASVs). For each combination in the grid, performance of the classifier on out-of-bag samples was assessed with 10-fold cross validation. Classifiers were then trained to predict each of the categorical factors (rootstock, scion, compartment, year, and site) along with all possible pairwise joint predictions. Tile plots were used to visualize output confusion matrix results. We determined relative importance of phyla in classification accuracy per factor, as well as ASVs that contributed considerably to classifier accuracy (i.e., high gini importance).

## Results

### Soil properties and soil microbiome showed site-specific differences

Soil texture, elemental composition and the diversity and richness of the soil microbiome differed across collection sites and by year. Soil texture was modeled in proportions of sand, silt, and clay, and clay was significantly different between collection sites (Figure 2A; Table S4; *P =* 0.041). Post-hoc comparisons revealed a 3.3% increase in clay content in Madera as compared to Merced (Figure 2A; *P =* 0.035). Sand and silt showed no significant differences across collection sites or years (Table S4). The principal component analysis of the elemental composition data showed clustering of samples by collection site (Figure 2B). PCs 1 and 2 collectively explained over 70% of the elemental variation (52.6% and 13.8% respectively), and each was with a linear model parameterized with collection site and year. Collection site showed a significant effect on the first PC (Table S5; *P* < 0.001) but not the second PC (Table S5), while collection year showed no significant effect on either PC. Post-hoc comparisons on the first PC model showed that the Madera site was differentiated from both Merced (*P* < 0.001) and San Joaquin (*P* < 0.001; Figure 1A) whereas the Merced vs San Joaquin comparison was non-significant.

**Figure 2.**
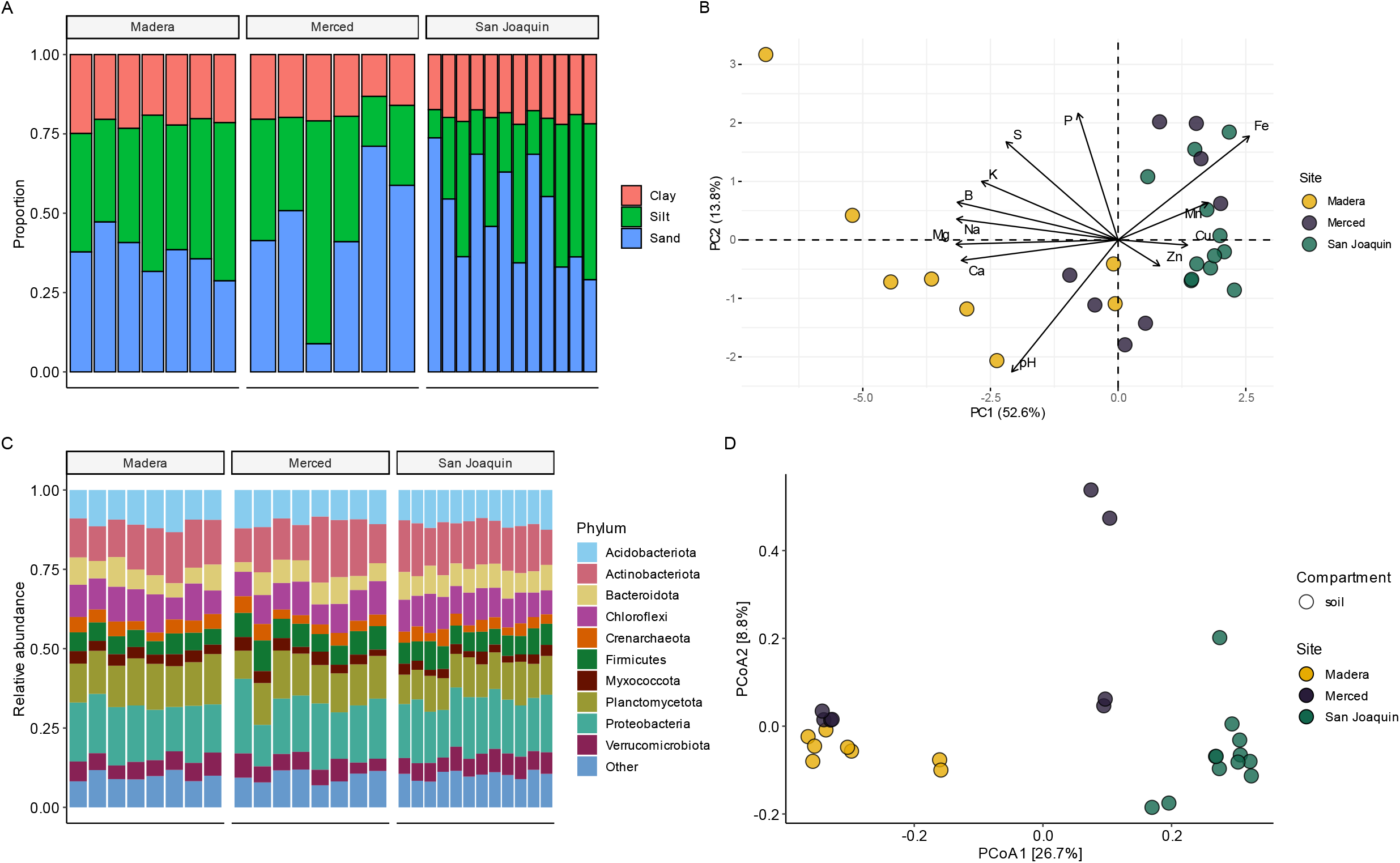
Soil properties and microbiome for the collection sites. **A)** The soil texture measured in proportions of sand, silt, and clay and **B)** a principal component analysis of soil elemental composition and pH, arrows represent loadings of elements and pH on principal components 1 and 2. **C)** Taxonomic barplots depicting the relative abundance of the top ten phyla for the soil microbiome and **D)** principal coordinate analysis on Bray-Curtis dissimilarity for soil samples. For both panels B and D, the colors correspond to collection sites in Figure 1A.

To assess the soil microbiome, we recovered 921,649 bacterial reads (n= 28; sample mean 32,916) across 8,838 ASVs after quality filtering. The top ten most abundant bacterial phyla recovered from soil samples were shared between sites (Figure 2C). Principal coordinate analysis of Bray-Curtis dissimilarity revealed clustering by site; we observed tight groupings for Madera and San Joaquin samples, but observed more variance in Merced samples, with some clustering closer to Madera samples (Figure 2D). PERMANOVA analysis further illustrated that differences in the composition of the soil microbiome were attributable to site (Table S6; *P* < 0.001), while sample collection year was non-significant. Both bacterial richness and diversity were impacted by collection year (Table S7; P = 0.0283 and P = 0.0386, respectively), with post-hoc comparisons showing that both richness and diversity were lower in 2018.

### Sugar content tracked vine development across the growing seasons

The sugar content of berries provided a standardized measure to assess development of the grapevines across season and site, as well as to characterize differences between scions ‘Cabernet Sauvignon’ and ‘Chardonnay’. Sugar content was responsive to the interaction of collection site, scion, and collection week (Table S8; *P* < 0.001). In the last collection week, ‘Cabernet Sauvignon’ was on average, 4.957 ± 0.824 °Brix lower in the northernmost collection site, San Joaquin, compared to °Brix observed in Madera and Merced (Figure 1A). All post-hoc comparisons for ‘Cabernet Sauvignon’ were significant (*P* < 0.001) with the exception of ‘Cabernet Sauvignon’ in the last collection week of 2018 (San Joaquin vs Merced; *P* = 0.108). For ‘Chardonnay’ we found a similar pattern, with the last collection week showing the most divergence in °Brix between for the sites, with San Joaquin on average 1.382 ± 0.813 lower than Madera and Merced (Figure 1A). Post-hoc comparisons for ‘Chardonnay’ showed that only the comparison of San Joaquin to Merced in the last collection week of 2018 was significant (*P* = 0.033). Given that berry sugar content tracks vine development across sites, season, and scions, we elected to use this measure in place of collection week in subsequent linear models.

### Bacterial diversity and richness were impacted by compartment, year, and rootstock by scion interactions

We generated bacterial sequence data for root (n=206), leaf (n=204), and berry (n=184) samples of grafted grapevines growing in three vineyards in the Central Valley of California. After quality filtering and removal of mitochondrial and chloroplast reads (2.2% and 1.9%, respectively), we recovered 28,993,602 reads across 45,332 ASVs. Decontam identified 183 ASVs as contaminants. Next, we filtered ASVs to retain only those present in five or more samples followed by read count filtering to remove samples with less than 1,000 reads. The resulting dataset comprised 594 samples with 7,981 ASVs and a total read count of 24,502,564 (sample mean 41,250). Finally, the ASV count matrix was normalized by applying a variance stabilizing transformation from the package DESeq2.

Plant compartment had the largest impact on Faith’s phylogenetic diversity index and richness (Table S9-10; Figure 3A). Root samples were significantly more diverse than both berries and leaves (Figure 3A; *P*<0.001) with mean values of 61.07, 4.63, and 5.71, respectively. Soil samples showed a slightly lower level of diversity than roots with a mean value of 52.08. In addition to compartment, Faith’s phylogenetic diversity index also varied significantly across collection years (Table S9; *P*<0.001), with samples collected in 2018 showing slightly higher diversity than samples collected in 2019. While site had a significant effect on Faith’s phylogenetic diversity index for plant compartments overall (Table S9; *P* = 0.041), no post-hoc comparisons were significant among sites. There was a significant interaction between rootstock and scion (Table S9; *P*<0.001), and post-hoc comparisons revealed that the rootstock ‘Teleki 5C’ grafted to the scion ‘Chardonnay’ had 6.91 lower mean bacterial diversity than other rootstocks (‘1103P’ vs ‘Teleki 5C’, *P*<0.001 and ‘Freedom’ vs ‘Teleki 5C’, *P*<0.001). This pattern was particular to the root samples (Figure 3B) and partially explained by a specific vineyard block at the Merced vineyard, Mer-4 (Table S1-2) featuring ‘Chardonnay’ grafted to ‘Teleki 5C’. Root samples within this block had a mean Faith’s phylogenetic diversity index of 37.10 ± 5.15, whereas the same rootstock/scion combination at the San Joaquin vineyard was 54.05 ± 2.39. Due to the experimental design, we are not able to fully disentangle the effects of particular vineyard blocks from rootstock/scion combinations. Although, the soil pH of this block was the lowest of the study at 5.7 (overall mean = 7.6; Table S2).

**Figure 3.**
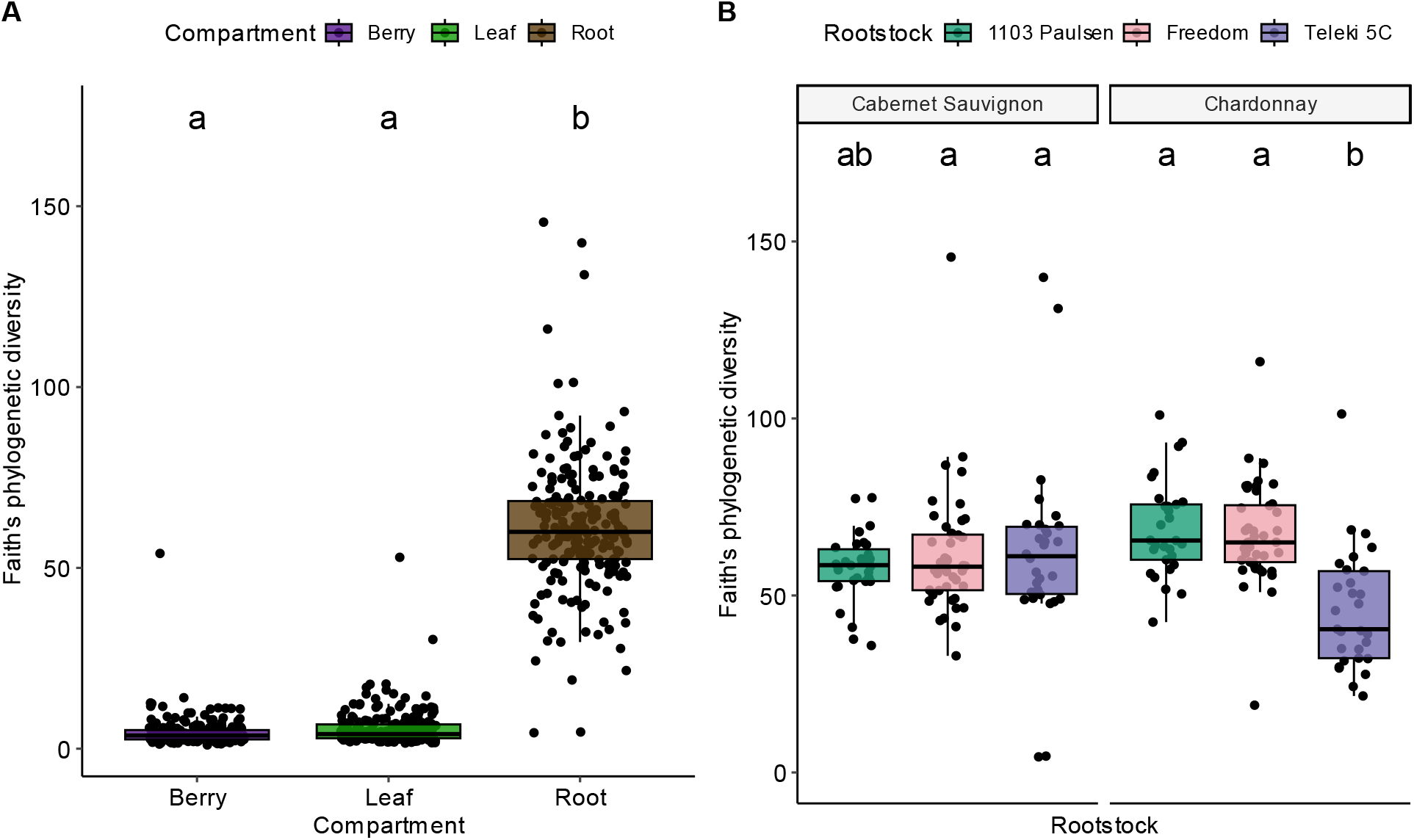
Faith’s phylogenetic diversity index was impacted by multiple factors of the experimental design. **A)** Plant compartment showed the largest effect on diversity and **B)** Root samples showed a rootstock by scion genotype effect on diversity. Boxplot hinges represent the 1^st^ and 3^rd^ quartiles with whiskers that represent 1.5 times the inter-quartile range; letters above box plots denote significant differences in means (TukeyHSD test).

### Bacterial composition of the root varied by site and by rootstock genotype but was relatively stable over time

Principal coordinate analysis on Bray-Curtis dissimilarity showed clear clustering of the bacterial composition of samples by plant compartment on axis 1 and 2 (Figure 4A), explaining 18.1% and 4.8% of the variance, respectively. The third axis, which explained 4.1% of the variance, showed clustering of root samples by the site of collection (Figure 4B), with San Joaquin separated from Madera and Merced. This pattern was unique to the root samples as berries and leaves were tightly clustered together on axis 3 (Figure 4B). The bacterial composition of the root compartment samples was relatively stable over the course of the growing seasons but was variable within a site, particularly Merced, and across the sites (Figure 4C; Table 2).

**Figure 4.**
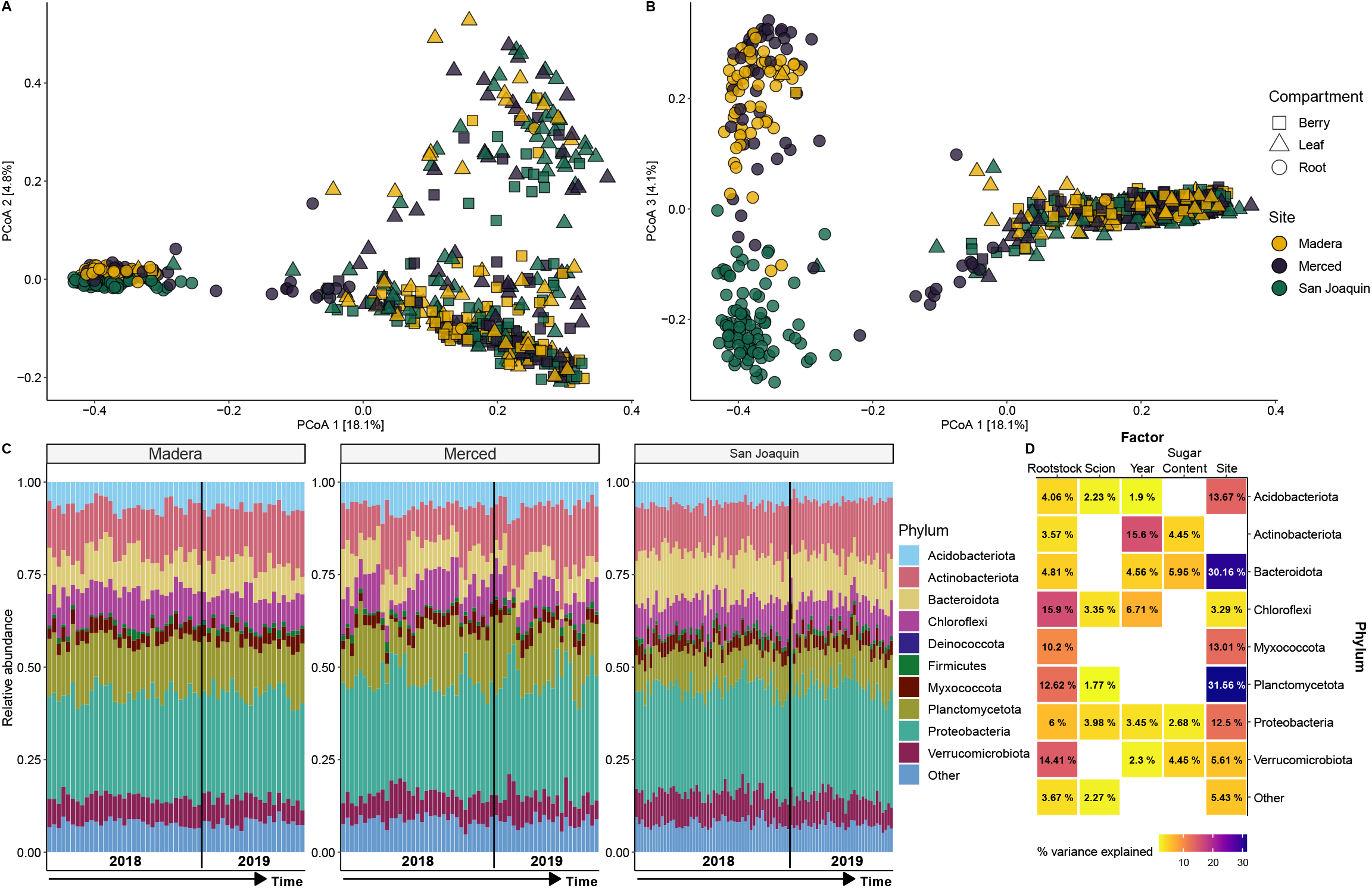
Principal coordinate analysis of Bray Curtis dissimilarity shows **A)** clustering of samples by compartment on axes 1 and 2 while **B)** axis 3 shows clustering of root samples by collection site. Samples are colored according to their collection site with shapes representing the three compartments sampled. Taxonomic barplots with the relative abundance of the top ten phyla for root samples **C)** delineated by collection site, each bar represents an individual sample and samples are ordered according to collection date with the black line denoting collection year. **D)** A tile plot summarizes the significant (*P*FDR<0.05) impact of the experimental factors on the relative abundance of the top ten phyla (Deinococcota and Firmicutes are omitted as no factors were significant).

**Table 2.** Permutational multivariate analysis of variance (PERMANOVA) using Bray-Curtis dissimilarity fit separately for each compartment.

| Factor | Berry |  | Leaf |  | Root |  |
| --- | --- | --- | --- | --- | --- | --- |
|  | R<sup>2</sup> | P | R<sup>2</sup> | P | R<sup>2</sup> | P |
| Rootstock | 0.011 | 0.474 | 0.010 | 0.258 | **0.046** | **0.001** |
| Scion | 0.007 | 0.110 | 0.005 | 0.280 | **0.018** | **0.001** |
| Sugar Content | **0.014** | **0.001** | **0.010** | **0.001** | **0.006** | **0.032** |
| Year | **0.015** | **0.001** | **0.034** | **0.001** | **0.014** | **0.001** |
| Site | 0.012 | 0.175 | 0.012 | 0.062 | **0.166** | **0.001** |
| Residual | 0.943 |  | 0.928 |  | 0.713 |  |

Bold values indicate significant results, $P < 0.05$ .

We examined the impact of experimental factors on the top ten phyla based off relative abundance for root samples (Figure 4D). Collection site impacted eight of the ten phyla and had the strongest impact overall on Planctomycetota (variation explained [VE] = 31.56%; *P*FDR<0.001), Bacteroidota (VE = 30.16%; *P*FDR<0.001), Acidobacteriota (VE = 13.67%; *P*FDR<0.001), Myxococcota (VE = 13.1%; *P*FDR<0.001), and Proteobacteria (VE = 12.5%; *P*FDR<0.001; Figure 4D). As expected, given the clustering in the PCoA (Figure 4B), the post-hoc comparisons of the other sites with San Joaquin were nearly all significant. Conversely, for Myxococcota and Proteobacteria, post-hoc comparisons between the other sites and Merced were non-significant. Only Deinococcota and Firmicutes, which together account for an average of 6.53% of relative abundance in roots samples, had no significant associations.

Next, we examined the influence of rootstock genotype and scion genotype on the relative abundance of the top ten phyla of the root compartment. Rootstock genotype impacted eight of the ten phyla, particularly Chloroflexi (VE = 15.9%; *P*FDR<0.001), Verrucomicrobiota (VE = 14.41%; *P*FDR<0.001), Planctomycetota (VE = 12.62%; *P*FDR<0.001), and Myxococcota (VE = 10.2%; *P*FDR<0.001; Figure 4D). Post-hoc comparisons showed that ‘Teleki 5C’ drove the differences, with higher relative abundances for Chloroflexi (mean = 1.9% increase), Planctomycetota (mean = 1.87% increase), and Myxococcota (mean = 0.57% increase) and lower relative abundance for Verrucomicrobiota (mean = 1.01% decrease). Compared to rootstock genotype, scion genotype had a smaller impact on relative abundance of the top ten phyla for roots. Only four of the ten phyla of the root compartment showed significant impacts for scion genotype. In Acidobacteria, Chloroflexi, Planctomycetota, and Proteobacteria, we found scion cultivar explained less than 5% of the variance (Figure 4D). Post-hoc comparisons showed that ‘Cabernet Sauvignon’ had a larger relative abundance of Acidobacteria, Chloroflexi, and Planctomycetota but the difference for each phylum was <1% between scion genotypes, while Proteobacteria had a smaller relative abundance (mean = 1.16% decrease).

Finally, we examined the influence of two aspects of time, collection year, and sugar content of the berries (an analog for vine development within a season) on the relative abundance of the top ten phyla of the root compartment (Figure 4D). Collection year significantly impacted six of ten phyla. Collection year had a particularly strong impact on the relative abundance of Actinobacteria (*PFDR*<0.001) explaining 15.6% of the variance (Figure 4D). The 2019 collection year showed enrichment of Actinobacteria in comparison to 2018 (mean = 2.73% increase). In comparison, sugar content was among the least impactful factors, with only four of the ten phyla showing significant impacts of this factor. Sugar content explained <6% of the variance for Actinobacteriota, Bacteroidota, Proteobacteria, and Verrucomicrobiota (Figure 4D).

### Bacterial composition of berries and leaves showed less predictable patterns across sites

Principal coordinate analysis showed clear clustering of bacterial communities by plant compartments (Figure 4A). Bacterial composition of the root showed a clear signature of site, we did not observe the same clustering in relation to site for berry and leaf samples (Figure 4A-B). For berries and leaves, PERMANOVA analysis showed a small effect of sugar content (Berries: R^2^ = 0.014, *P* = 0.001; Leaves R^2^ = 0.010, *P* = 0.001) and collection year on Bray-Curtis dissimilarity (Berries: R^2^ = 0.015, *P* = 0.001; Leaves R^2^ = 0.034, *P* = 0.001; Table 2). Taxonomic barplots of the top ten phyla of shoot compartments showed fluctuations in relative abundance but were non-specific to collection sites (Figure S4). Using the same linear model framework as above, we found that, for berries, only a single phylum, Bacteroidota showed a significant effect of collection year (VE = 4.12%: *P*FDR = 0.029). Post-hoc comparisons revealed that berries in 2018 had on average 2% greater abundance of Bacteroidota. For leaves, we found only a couple phyla were impacted by the experimental factors. First, Firmicutes showed patterning by collection site (VE = 6.00%; *P*FDR = 0.010), post-hoc comparisons of Firmicutes showed that the Merced site had, on average, 4.29% increased relative abundance compared to the other two sites (Figure 1A) for leaf samples. Of note, Firmicutes did not show any significant impact of the experimental design for root samples (Figure 4D). Second, Actinobacteriota showed patterning by the collection year (VE = 7.02%; *P*FDR < 0.001), post-hoc comparisons of Actinobacteria showed a 5.42% decreased in relative abundance for 2018 vs 2019.

### Machine learning accurately predicts the root compartment but not berries or leaves

We used machine learning to identify the factors that were predictable across the experimental design (Figure S5) and to pinpoint the ASVs that aided the accuracy of those predictions (Figure S6-7). Overall, model accuracy was 68% (Figure S5A); however, classifier was near perfect when predicting root samples (F1 = 0.986; Table S11) but was less accurate when predicting leaf (F1 = 0.525) and berry (F1 = 0.565). For rootstock genotype, model accuracy was 50% (Figure S5B) and ‘Freedom’ was the most predictable genotype (F1 = 0.623; Table S11) while ‘1103 Paulsen’ and ‘Teleki 5C’ were considerably lower (F1 = 0.431 and 0.204, respectively). Model accuracy for collection site was 59% (Figure S5C), the most predictable site was San Joaquin (F1 = 0.672), followed by Merced (F1 = 0.559) and then Madera (F1 = 0.431; Table S11). For collection year model accuracy was 83% (Figure S5D) and an F1 score of 0.882 (Table S12). For scion genotype, model accuracy was 61% (Figure S5E) and with an F1 score of 0.623 (Table S12). Finally, for sugar content, model wide accuracy was 71% (Figure S5F) and an F1 score of 0.795 (positive class was pre-ripening; Table S12).

When examining the phyla that were the most important in the classifier’s predictions, we found that across all factors many of the phyla have similar relative importance to their respective classifier (Figure S6). Proteobacteria was the most important phylum making between 39.7-54.6% of the relative importance to the classifier and Actinobacteria was the second most important phylum with between 14-24.9% of the relative importance to the classifier (Figure S6). Crenarchaeota was only important to the classification of plant compartments and collection sites with all ASVs showing affinity to the root compartment and some showing site-specificity. ASVs of this phylum (Figure S7), were annotated to the family Nitrososphaeraceae of which isolated members are known to oxidize ammonia and have been recovered previously from soils (Tourna *et al*. 2011; Stieglmeier *et al*. 2014).

## Discussion

The goal of this study was to investigate factors influencing bacterial communities of grapevine roots, leaves, and berries, including rootstock genotype, scion genotype, and vineyard site, within growing seasons and over multiple years. We observed differences in soil texture, elemental composition, and bacterial communities among vineyard sites sampled in this study. We detected differences in bacterial composition of grapevine root compartments across sites; however, site specific differences were less pronounced in the microbiota of the berries and leaves. Both rootstock and scion genotype impacted composition and diversity of vine microbiota. Using brix (berry sugar content) as a proxy for development, we observed only minor associations between developmental stage and bacterial community composition of the berries and leaves. This suggests that berry and leaf bacterial communities are likely seeded from tissues with mature microbiomes and undergo largely stochastic changes in community composition across the season.

### Soils and the soil microbiome differ across sites

In the cultivation of wine grapes, the term terroir describes regional environmental factors, including soil properties, geography, and climate, that influence characteristics of wine (Seguin 1986; Van Leeuwen and Seguin 2006; Van Leeuwen *et al*. 2018). These same factors influence the soil microbiome, a primary reservoir of microorganisms that colonize and ultimately become associated with plants (Chi *et al*. 2005; Lundberg *et al*. 2012; Bulgarelli *et al*. 2012; Zarraonaindia *et al*. 2015; Liu *et al*. 2017). Thus, many studies have uncovered a role for microorganisms in shaping terroir of wine, often called the microbial terroir (Bokulich *et al*. 2014; Gilbert *et al*. 2014; Knight *et al*. 2015). Previous research has shown that across regional scales, grapevine musts (crushed berry clusters) and wines exhibit site specific microbiota (Bokulich *et al*. 2014; Setati *et al*. 2015; Garofalo *et al*. 2016; Jara *et al*. 2016), which associated with the metabolomic composition of the wine (Bokulich *et al*. 2016; Zhou *et al*. 2021).

In this study, we demonstrated that vineyard soils within the Central Valley of California varied in elemental composition and soil texture (Figure 2B&D). Further, differences in soil microbial communities were apparent across vineyards, an observation consistent with previous work in other viticultural regions (Zarraonaindia *et al*. 2015; Burns *et al*. 2015; Coller *et al*. 2019; Zhou *et al*. 2021). For instance, a recent global survey of vineyard soils across five continents (Gobbi *et al*. 2022), revealed prokaryotic communities bear a signature of spatial distance, with dissimilarity strongly positively correlated with geographic scales (regional to continental). Here, we found similar patterns at an intra-regional scale (∼177 kilometers apart between farthest vineyards) in the Central Valley of California. This work contributes to a growing of literature documenting site-specific differences in biotic and abiotic properties of vineyard soils.

### Vineyard site influences bacterial composition of the grapevine root, but not the shoot

Within grapevines, bacterial composition bears a strong signature of compartment with roots, leaves, and berries exhibiting distinct microbiota (Zarraonaindia *et al*. 2015; Deyett and Rolshausen 2020; Swift *et al*. 2021). Data presented here confirm that compartment specific dynamics primarily dictate the grapevine microbiota, an observation consistent with previous studies across plant species (Coleman-Derr *et al*. 2016; Brown *et al*. 2020). Thus, we might expect that grapevine compartments respond differently to changes in the soil microbiome. For example, Zarraonaindia *et al*. (2015) found that bacterial communities of the soil differed across several vineyards in the Northeastern United States, but those differences were not reflected in plant compartments. Conversely, a study conducted across the California coastal growing regions showed regional patterning for grapevine musts (crushed berries) for both bacterial and fungal communities (Bokulich *et al*. 2014). The amount of variation explained by region and vineyard for bacteria was generally lower than for fungi, indicating that kingdoms exhibit different diversity patterns across local and regional environmental conditions (Gobbi *et al*. 2022).

Data presented here indicate that the bacterial community of the root was strongly influenced by vineyard site (Table 2; Figure 3B&D), whereas bacterial communities of leaves and berries did not show site-specific patterning (Table 2; Figure S4). Differences in how collection site impacts grapevine compartment bacterial composition may reflect various biological and environmental factors. From a biological perspective, the root system is the only compartment assessed here that is in direct, sustained contact with the soil. Thus, it makes sense that the bacterial community of the root would bear a stronger signature of the soil microbiome than the bacterial communities of the above-ground compartments of the vine, which were not in direct contact with the soil. While it has been consistently shown that majority of the microbiota of the vine enters through the roots (Zarraonaindia *et al*. 2015; Swift *et al*. 2021). It is possible that filtering occurs, across the graft junction, by the scion genotype, as well as through selective pressure in the leaf and berry compartments, which may serve to dilute the signal of the site from the roots to the shoot system.

Beyond the biology of the soil-root connections, there may be other environmental factors that dictate compositional patterns in bacterial communities of above-ground vs. below-ground compartments (Allard *et al*. 2020; Mechan Llontop *et al*. 2021; Morales Moreira *et al*. 2021). For example, rainfall serves as a dispersal mechanism for microorganisms, either through deposition from the atmosphere via precipitation (Woo *et al*. 2018; Cáliz *et al*. 2018; Aho *et al*. 2020; Mechan Llontop *et al*. 2021) or through water splashes containing soil (Cevallos-Cevallos *et al*. 2012; Monaghan and Hutchison 2012; Allard *et al*. 2020). Rainfall events catalyzed community succession in canola (*Brassica napus* L.) leaf microbiota (Copeland *et al*. 2015). In both 2018 and 2019, vineyards sampled for this study did not experience any precipitation during the sampling window and all hydration for the vine was supplied by drip irrigation (weather.gov). Consequently, it is unlikely the soil microbes would be dispersed to aerial parts of the plant (leaves and berries) via water splashes at these vineyards during our sampling window.

A second possible environmental factor underlying our observation that root microbiota bears a signature of the surrounding soil, but leaves and berries do not, is the influence of the wind. Wind dispersal of soil microbiomes as observed in other plant systems (Rastogi *et al*. 2012; Maignien *et al*. 2014; Ottesen *et al*. 2016) is a more likely factor in the Central Valley of California, as the majority of vines sampled in this study were covered in soil. A recent study found that airborne fungal microbiota (collected as settled dust) across the Central Valley of California were distinct from coinciding soil fungal microbiota, exhibiting remarkable similarity even from sites over 150 km apart (Wagner *et al*. 2022). Similarly, leaf and berry microbiota described here, despite variance in soil and root microbiota, are largely homogeneous, possibly due to airborne dispersal and deposition of dust/soil particles and the absence of physical disturbance by rain.

### The influence of rootstock and scion genotype

Beyond the influence of the vineyard environment, there is evidence to suggest that both the rootstock and scion genotype play a role in shaping the bacterial communities in vine compartments. We detected an effect of both rootstock and scion genotype on bacterial composition of the root (Figure 4; Table 2), as well as the bacterial diversity of the vine as a whole (Table S9-10). Interestingly, while the bacterial composition of root samples showed a signal of both rootstock and scion genotype, berry and leaf bacterial composition showed no significant patterning by either host genotype (Table 2). This is consistent with a recent study, where rootstock by scion interactions were associated with changes in the diversity and composition of root bacterial communities in grapevine (Marasco *et al*. 2022) and previous studies examining rootstock-specific rhizospheres of multiple rootstock genotypes (D’Amico *et al*. 2018; Marasco *et al*. 2018; Berlanas *et al*. 2019). The phenotype most likely to drive host genotypic differences in vine bacterial composition is the root exudate profile, consisting of sugars, organic acids, and amino acids, among other compounds (Sasse *et al*. 2018). In *Arabidopsis thaliana*, root exudate variation was linked to the genomic variation among *A. thaliana* accessions (Mönchgesang *et al*. 2016) and has been associated with distinct rhizosphere bacterial communities (Micallef *et al*. 2009).

Rootstocks studied here, and employed in most commercial vineyards, are of complex genetic backgrounds with two genetically distinct species serving as parents (Table 1; Riaz *et al*. 2019). Thus, there is likely substantial variation between the rootstock genotypes that may influence root exudate profiles and, subsequently, the associated root microbiota. Relative to the rootstock genotype, the genotype of the grafted scion had a smaller overall influence on the composition of the root microbiome. This could be explained by proximity, with the rootstock genotype indirect contact with soil microorganisms. Further, relative to rootstock genotypes, the scion genotypes are more genetically similar to one another because both are cultivars of a single domesticated species, *Vitis vinifera*. Despite this, we found evidence for rootstock by scion interactions indicating both genotypes contribute to the formation of the root microbiome. Experiments utilizing reciprocal rootstock/scion combinations can begin to quantify the contribution of both the rootstock and scion in shaping the grapevine microbiome.

### Bacterial diversity is stable over the growing season and may be decoupled from patterns of fungal diversity

By sampling across the growing season over multiple years, our experimental design allowed us to investigate the progression of bacterial communities of multiple compartments of the vine. We used berry sugar content to track vine development across the season and determined that the bacterial diversity of grapevine compartments generally do not shift based on vine developmental stage (Table S9 & S10). Despite this, we observed compartment specific seasonal shifts in bacterial composition, however, sugar content and year explained only a small proportion of the variance (Table 2; Figure 4C & S4). Given that our experimental factors only captured a small amount of variation in the bacterial composition of both berries and leaves, this may indicate that microbiota of these aerial compartments are influenced primarily by macro- and microclimatic conditions rather than seasonal or developmental patterns.

Data presented here focus on the bacterial communities of the grapevines. It is important to note that results for bacteria presented here and elsewhere (Pinto *et al*. 2014), contrast with some patterns of fungal communities, where plant development has been shown to be a stronger predictor of shoot system community composition (Liu and Howell 2021). Other studies examining the microbiota of the berry epidermis reported associations between development stages and fungal (Zhu *et al*. 2021; Ding *et al*. 2021) and bacterial communities (Wei *et al*. 2022). Given the sample size here (berries n=184; leaves n=204), we should be able to detect even subtle changes in community diversity and composition across vine development. However, it is possible that the sampling window may have been too narrow. Including samples from the earliest developmental stages (leaf flush and fruit set) to latest stages (senescence and harvest), may have resulted in a stronger signature of development on the microbiota.

## Conclusions

Uncovering factors shaping plant-associated microbiomes from annuals to long-lived perennials holds potential for numerous applications in agriculture. Work described here demonstrates site-specific biogeographical patterns in bacterial communities in the soil, as well as corresponding patterns of in the root compartments of grafted grapevines. However, bacterial communities of berries and leaves reflected patterning by development and collection year, not site. This result provides further resolution into the processes underlying microbial terroir in grafted grapevines, illustrating that different compartments are uniquely impacted by microbial biogeography. In addition, we show that rootstock genotype more strongly influences the bacterial communities of the compartments than the scion genotype. Future studies should leverage experimental designs that allow simultaneous sampling of multiple plant compartments across both space and time along with metagenomic techniques to probe the functional significance of differences between microbiomes.

## Acknowledgements

We thank Peter Cousins (E. & J. Gallo Winery) for logistical support through all phases of collections and vineyard managers and staff Amanpreet Virk, Danielle Hopkins, Jorge Fuentes, Keith Striegler, Lindsay Jordan, and Octavio Viveros for their support. In addition, we are grateful to collaborators Mani Awale (University of Missouri), Dan Chitwood (Michigan State University), Misha Kwasniewski (University of Missouri), Laura Klein (Saint Louis University), and Matt Maimaitiyiming (University of Missouri) for assistance in the field. This project was made possible in part by funds supporting a summer training program in which undergraduate students from partnering institutions contributed to field work in commercial vineyards in California. We acknowledge the following students for their help in this work: Julie Curless (Missouri State University), Zach Helget (South Dakota State University), Anh Ly (Missouri State University), Dalton Gillig (University of Missouri), Vy Nguyen (Missouri State University), Leah Pinkner (Missouri State University), and Karoline Woodhouse (South Dakota State University). This work was funded by the National Science Foundation Plant Genome Research Program 1546869. JFS was supported by an NSF Graduate Research Fellowship under Grant No. 1758713 and the Saint Louis University graduate program. Lastly, we would like to thank members of the Miller and Wagner lab for helpful comments in preparing this article.

## Data availability

All raw sequencing data is available on NCBI SRA under BioProject ID PRJNA849941 and SRA accessions SRR19735936-SRR19736586. All code to reproduce the analysis and figures is available on GitHub at https://github.com/Kenizzer/California_Transect_Microbiome.

## Author Contributions

JFS, ZM, and AJM designed the experimental and collection design. JFS and ZM performed sample collection. JFS and GET performed sample processing. JFS and ZM performed data analysis. All authors contributed to data interpretation, the writing and reviewing of the manuscript, and approved the final draft.

## Supplemental Figures

**Figure S1.**
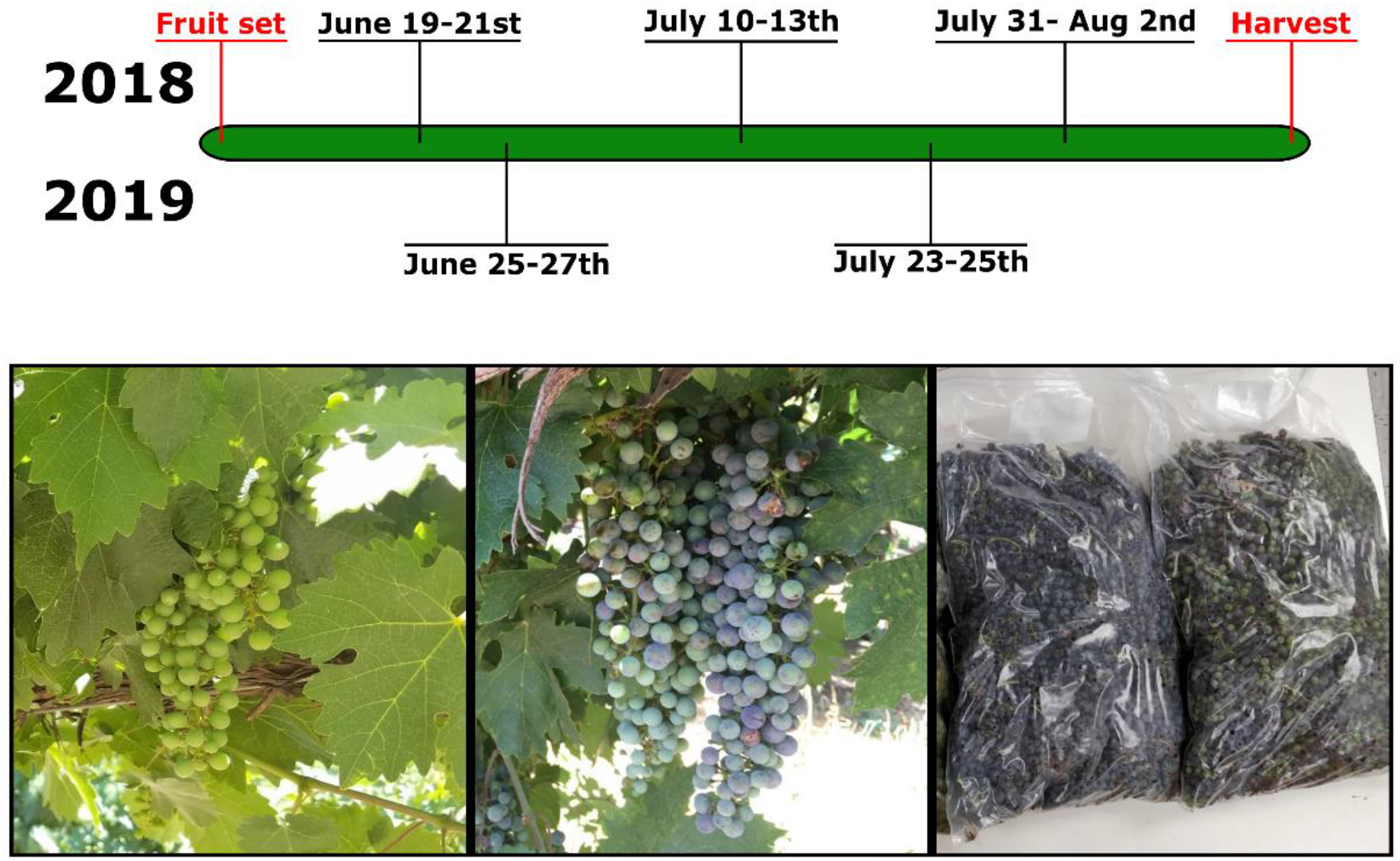
Collections were staggered across the growing season to capture the development of the vines. Photos display the progress of the vines for collections in 2018.

**Figure S2.**
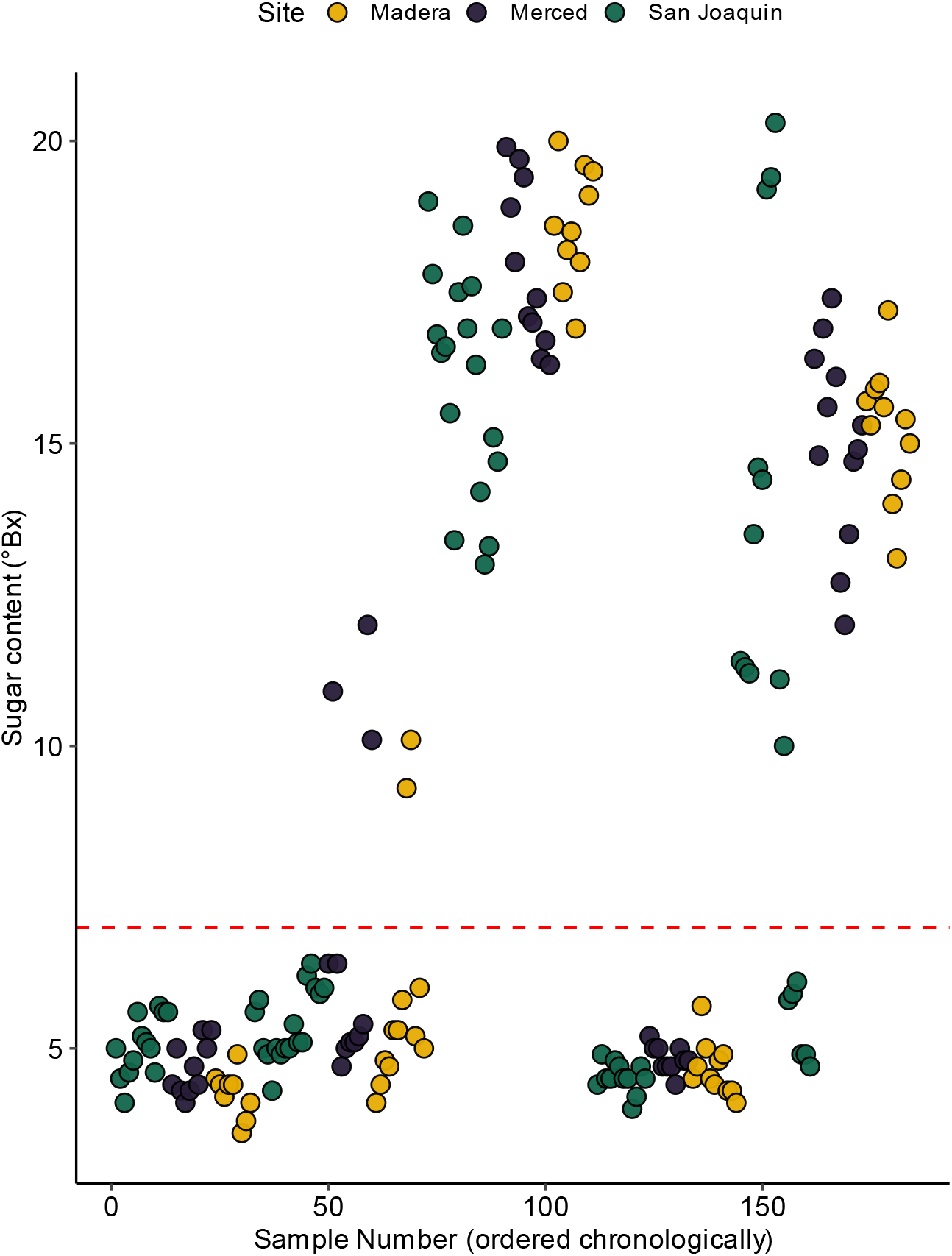
Discretization of sugar content values. The horizontal dashed line denotes where sugar content values (measured in °Bx) for samples were split in order to discretize for statistical analysis, values 3-7°Bx were labeled pre-ripening and values >7°Bx were labeled ripening. Points are colored according to the collection site.

**Figure S3.**
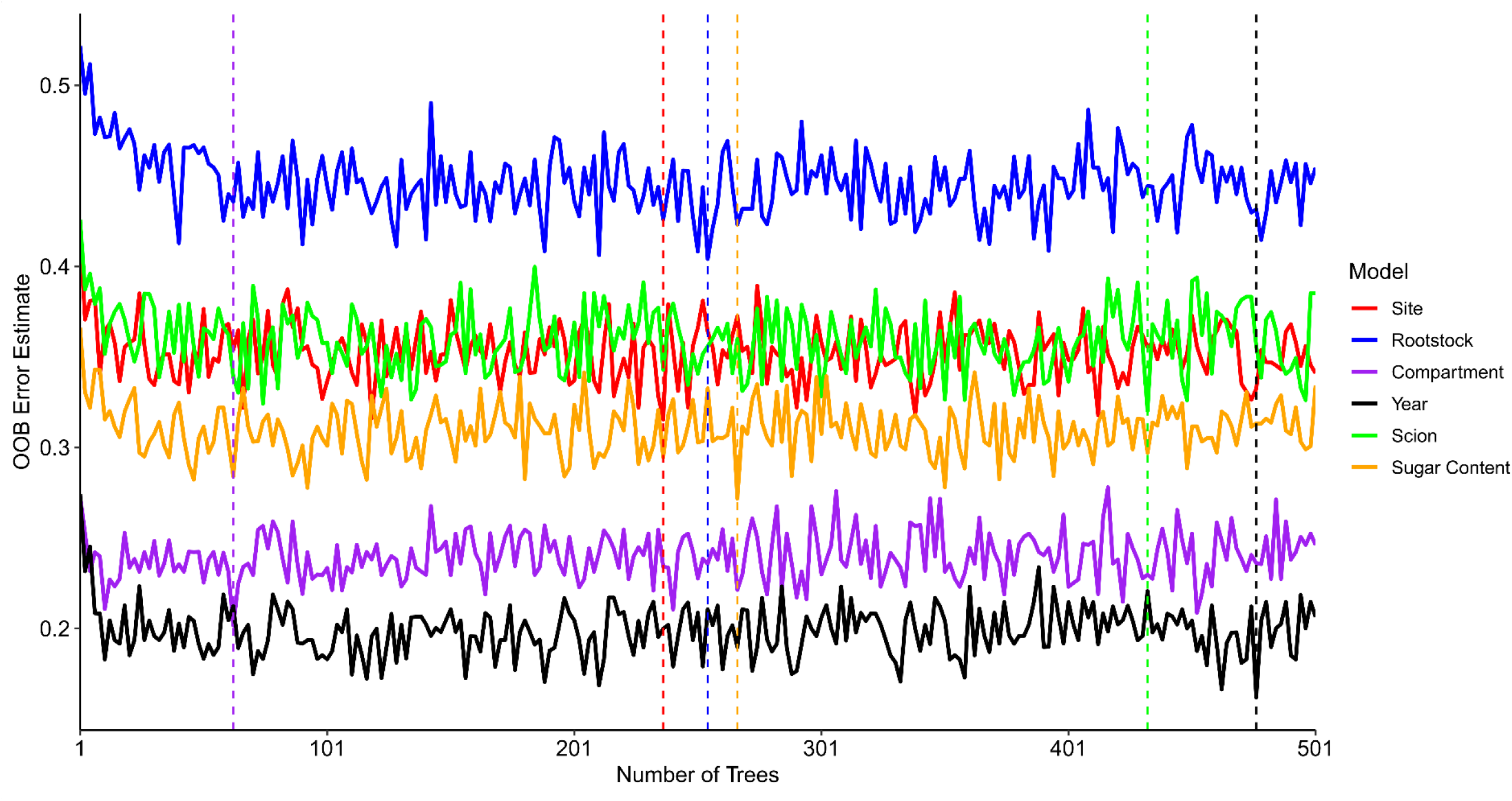
Out-of-bag error estimates for random forest models, across values for the number of trees, attempting to predict collection site, rootstock genotype, plant compartment, collection year, scion genotype, and sugar content. Dashed lines represent the minimum error estimate returned per model.

**Figure S4.**
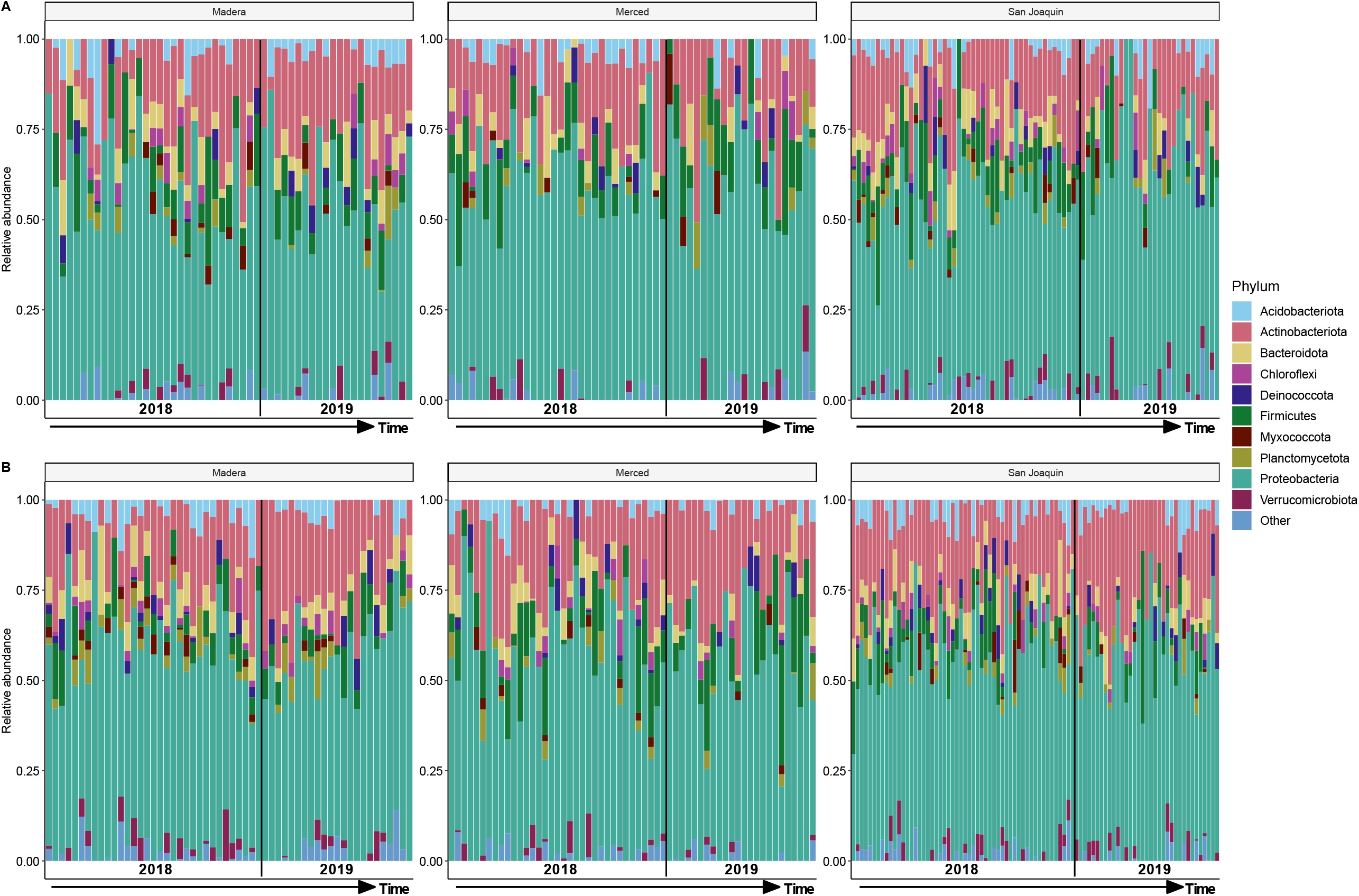
Taxonomic barplots with the relative abundance of the top ten phyla for A) berry samples and B) leaf samples delineated by collection site, each bar represents an individual sample and samples are ordered according to collection date with the black line denoting collection year. All phyla below the top ten are condensed into the other category.

**Figure S5.**
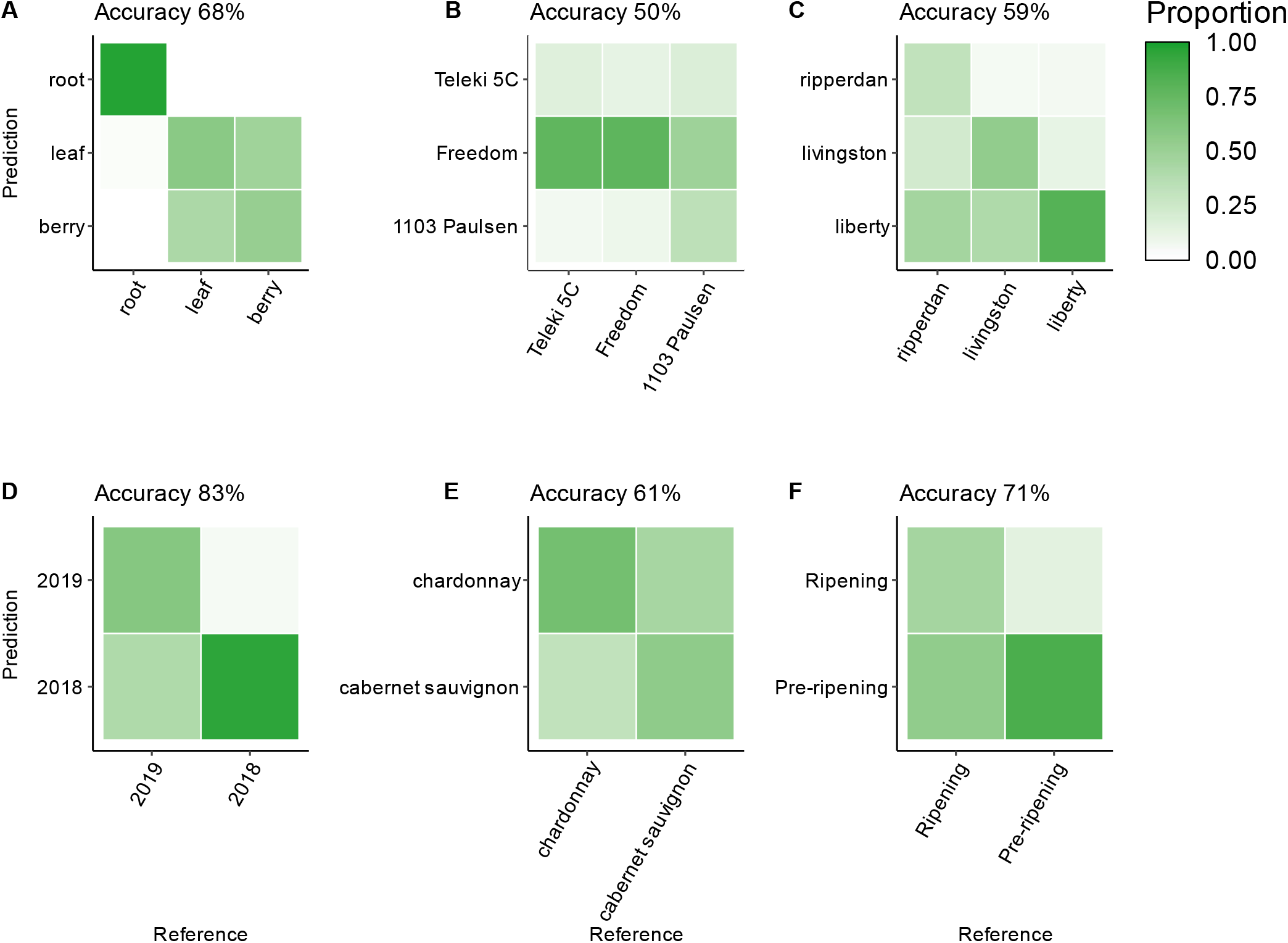
Confusion matrices for each experimental factor; A) plant compartment, B) Rootstock genotype, C) Scion genotype, D) Collection site, and E) Collection year. The model was trained using an 80:20 data split (80% train, 20% test) with 10-fold cross validation, shading represents the proportion of predictions for a label. Labels on the left represent the predicted labels and the labels on the bottom represent the actual labels. The model wide accuracy is given above each confusion matrix, class-wise statistics are provided in table S11-12.

**Figure S6.**
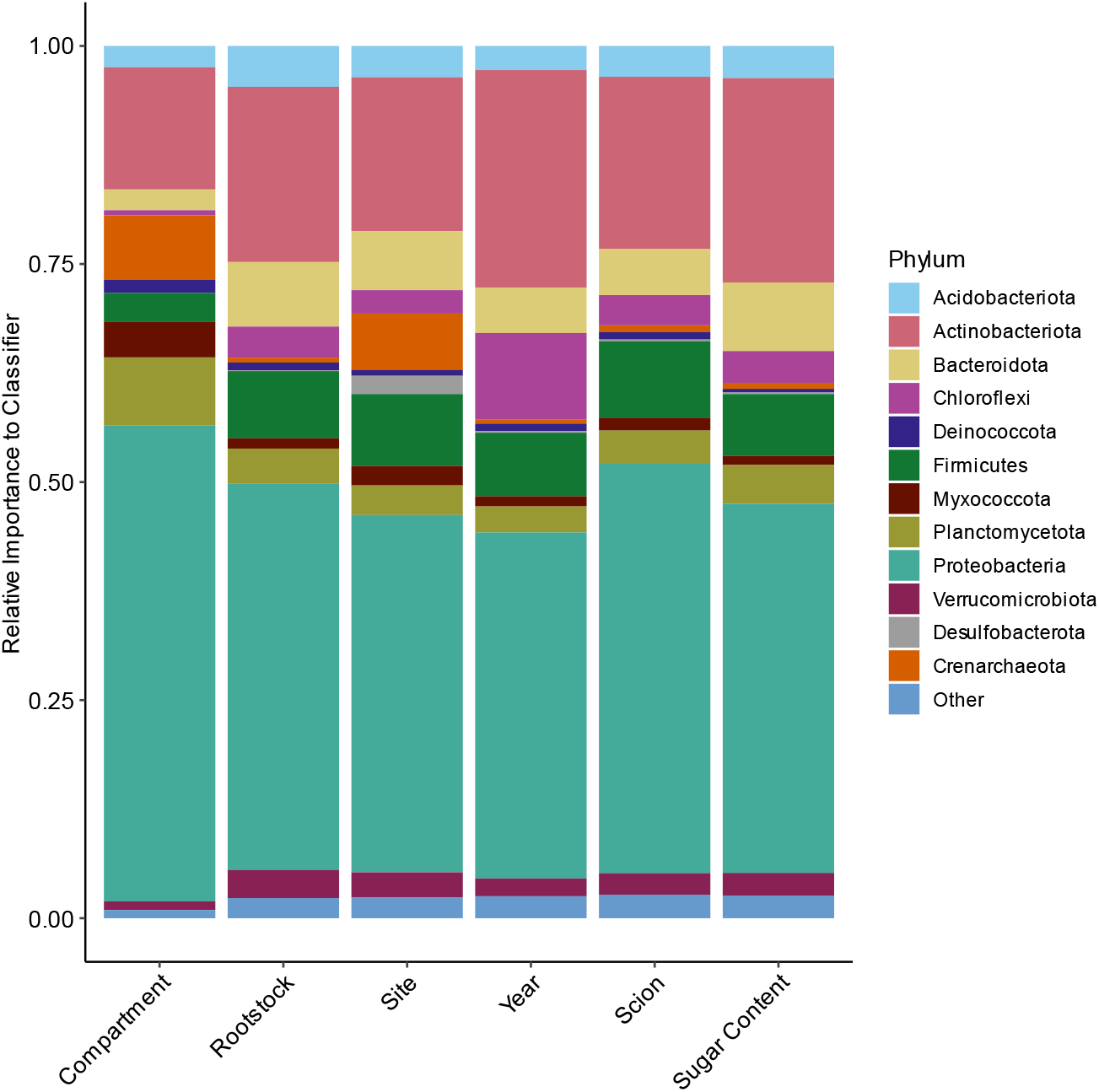
Relative importance of phyla to random forest classifiers for each experimental factor.

**Figure S7.**
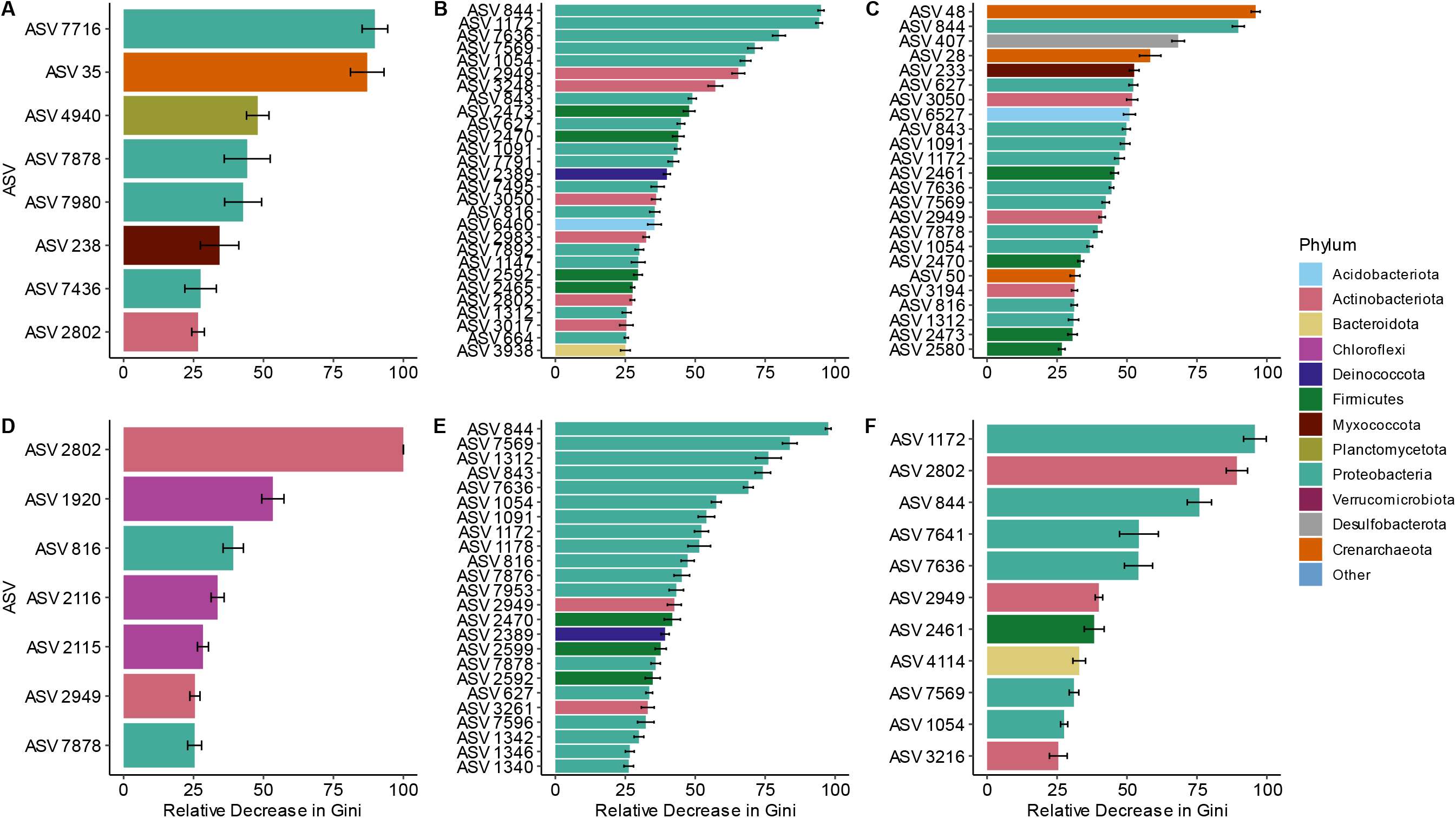
Individual ASVs that contribute to the random forest classifiers trained for; A) plant compartment, B) rootstock genotype, C) collection site, D) Collection year, E) scion genotype, and F) Sugar Content. Gini importance was assessed for all ASVs on the out-of-bag samples for each fold of the cross validation (*n*=10) and was scaled from 0 to 100 (higher being more important to the trained classifier). Only ASVs showed a greater than 25% decrease in gini importance are shown.

**Table S1.** Characteristics of the vineyard blocks used within the study. “Cab. Sauv.” = Cabernet Sauvignon

| Block | County | Scion | Rootstock | Est Year | Acreage | Vine Spacing | Trellis System | Orientation | Cover Crop |
| --- | --- | --- | --- | --- | --- | --- | --- | --- | --- |
| Mad-1 | Madera | Cab. Sauv. | 5C | 1995 | 42.11 | 7x11 | Horizontally divided double curtain | N/S | *Poaceae* spp. |
| Mad-2 | Madera | Cab. Sauv. | Freedom | 1995 | 37.89 | 7x11 | Simple curtain | N/S | *Poaceae* spp. |
| Mad-3 | Madera | Chardonnay | 1103P | 2009 | 76.45 | 6x9 | Vertically divided double curtain | E/W | *Poaceae* spp. |
| Mad-4 | Madera | Chardonnay | Freedom | 2008 | 37.64 | 6x9 | Vertically divided double curtain | E/W | *Poaceae* spp. |
| Mer-1 | Merced | Cab. Sauv. | Freedom | 1996 | 185 | 5x11 | Horizontally divided double curtain | E/W | *Poaceae* spp. |
| Mer-2 | Merced | Cab. Sauv. | 1103P | 2012 | 111.9 | 6x9 | Simple curtain | E/W | *Poaceae* spp. |
| Mer-3 | Merced | Chardonnay | Freedom | 2010 | 18.8 | 6x9 | Vertically divided double curtain | E/W | *Poaceae* spp. |
| Mer-4 | Merced | Chardonnay | 5C | 1995 | 176.8 | 4x10 | Horizontally divided double curtain | E/W | *Poaceae* spp. |
| San-1 | San Joaquin | Chardonnay | Freedom, 1103P, 5C | 1994 | 0.8 | 7x10 | Simple curtain | N/S | N/A |
| San-2 | San Joaquin | Cab. Sauv. | Freedom, 1103P, 5C | 1994 | 0.8 | 7x10 | Simple curtain | N/S | N/A |

**Table S2.**
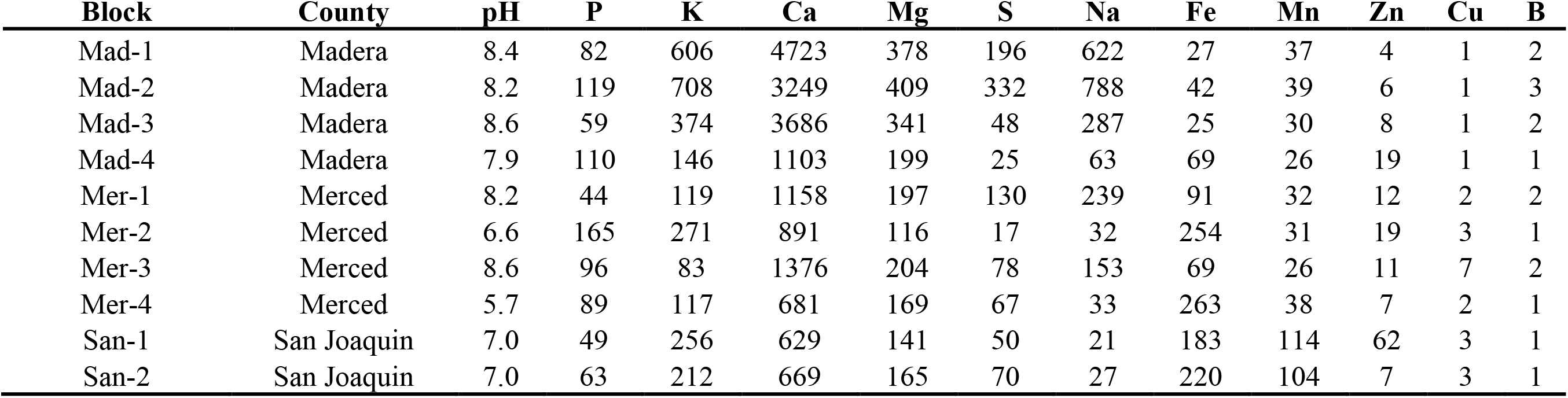
Soil elemental composition of the vineyard blocks used within the study. Values are reported in mg/kg (ppm) with the exception of pH which is reported in the standard scale.

**Table S3.** Optimal hyperparameters for training machine learning models.

| Model | mtry | Minimum node size | Number of trees |
| --- | --- | --- | --- |
| Rootstock | 1606 | 10 | 255 |
| Compartment | 1606 | 10 | 63 |
| Site | 808 | 5 | 237 |
| Year | 1606 | 10 | 477 |
| Scion | 4000 | 5 | 433 |
| Sugar Content | 5596 | 10 | 267 |

**Table S4.**
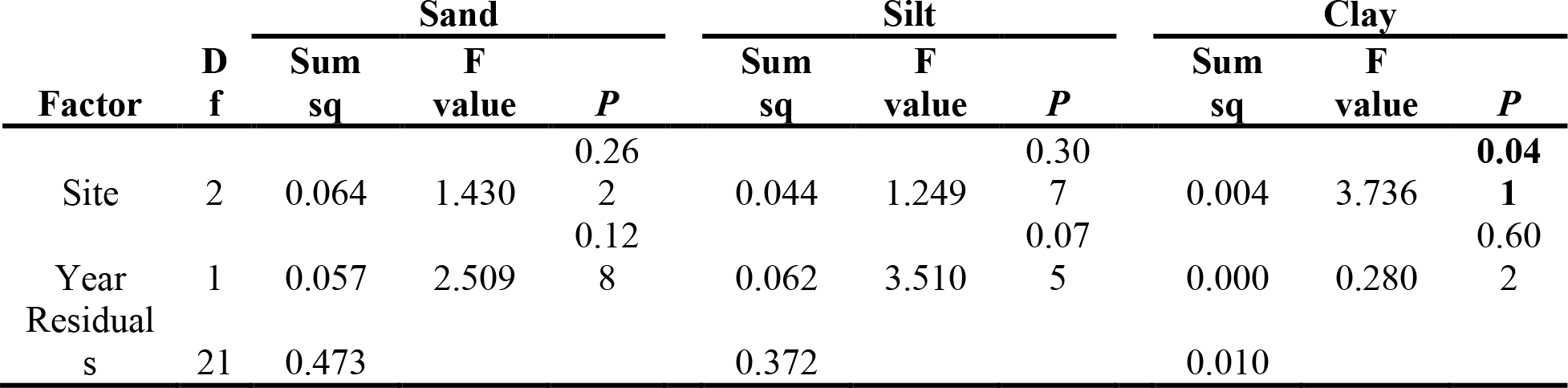
Linear model results for soil texture. Type-2 ANOVA table.

**Table S5.**
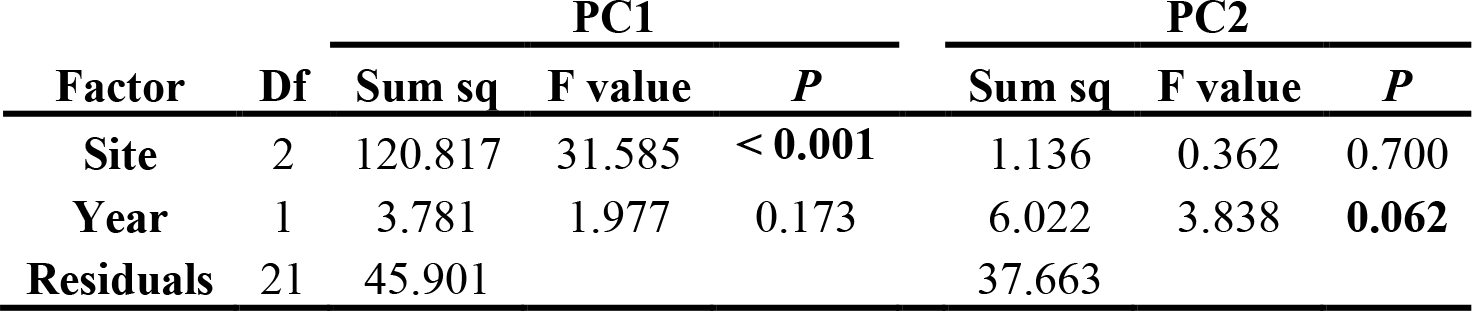
Linear model results for soil elemental composition principal components (PC). Type-2 ANOVA table.

| Factor | Df | PC1 |  |  | PC2 |  |  |
| --- | --- | --- | --- | --- | --- | --- | --- |
|  |  | Sum sq | F value | P | Sum sq | F value | P |
| Site | 2 | 120.817 | 31.585 | **< 0.001** | 1.136 | 0.362 | 0.700 |
| Year | 1 | 3.781 | 1.977 | 0.173 | 6.022 | 3.838 | **0.062** |
| Residuals | 21 | 45.901 |  |  | 37.663 |  |  |

**Table S6.**
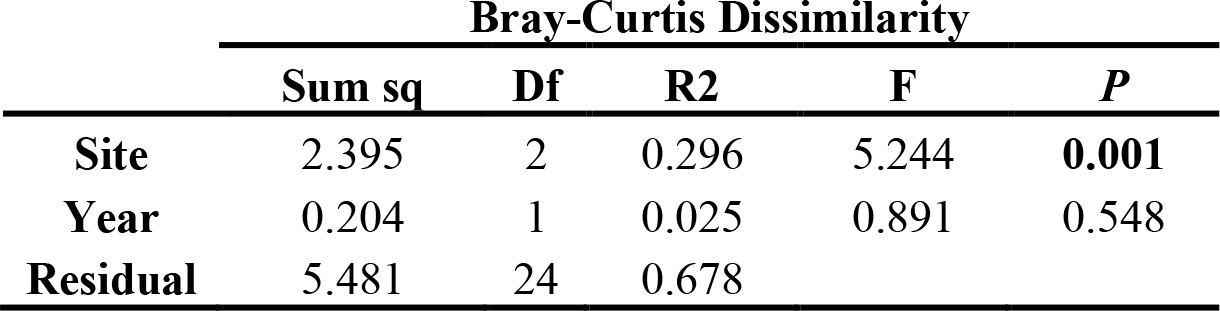
Linear model results for Bray-Curtis Dissimilarity for soil samples. Type-2 ANOVA table.

**Table S7.** Linear model results for alpha diversity statistics for soil samples, Chao1 index and Faith’s phylogenetic diversity. Type-2 ANOVA table.

| Factor | Df | Chao1 |  |  | Faith's phylogenetic |  |  |
| --- | --- | --- | --- | --- | --- | --- | --- |
|  |  | Sum sq | F value | *P* | Sum sq | F value | *P* |
| **Site** | 2 | 13082 | 0.681 | 0.515 | 43.49 | 0.464 | 0.634 |
| **Year** | 1 | 52231 | 5.441 | **0.028** | 224.64 | 4.792 | **0.039** |
| **Residuals** | 21 | 230375 |  |  | 1125.07 |  |  |

**Table S8.**
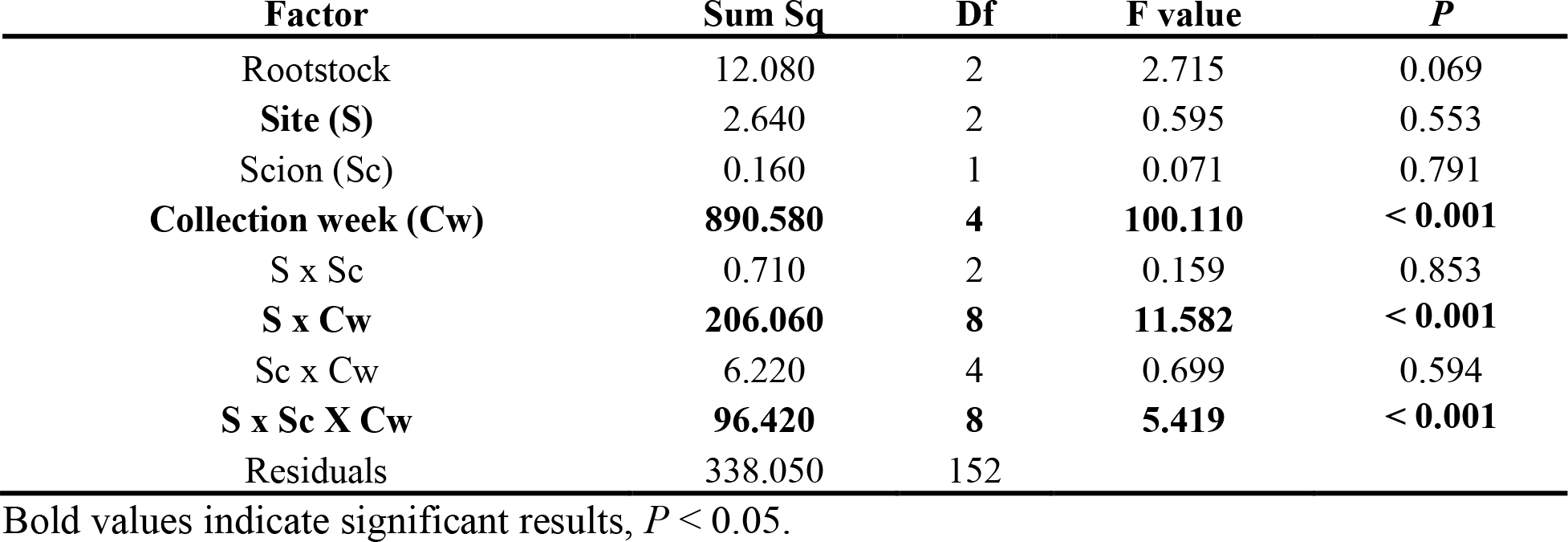
Linear model results for sugar content of the berries. Type-3 ANOVA table.

| Factor | Sum Sq | Df | F value | *P* |
| --- | --- | --- | --- | --- |
| Rootstock | 12.080 | 2 | 2.715 | 0.069 |
| **Site (S)** | 2.640 | 2 | 0.595 | 0.553 |
| Scion (Sc) | 0.160 | 1 | 0.071 | 0.791 |
| **Collection week (Cw)** | **890.580** | **4** | **100.110** | **< 0.001** |
| S x Sc | 0.710 | 2 | 0.159 | 0.853 |
| **S x Cw** | **206.060** | **8** | **11.582** | **< 0.001** |
| Sc x Cw | 6.220 | 4 | 0.699 | 0.594 |
| **S x Sc X Cw** | **96.420** | **8** | **5.419** | **< 0.001** |
| Residuals | 338.050 | 152 |  |  |

Bold values indicate significant results, $P < 0.05$ .

**Table S9.** Linear mixed model results for Faith’s Diversity index. Type-3 ANOVA table..

| Factor | Sum Sq | Mean Sq | F value | P |
| --- | --- | --- | --- | --- |
| **Rootstock** | **42.500** | **21.230** | **3.475** | **0.034** |
| Scion | 0.500 | 0.500 | 0.082 | 0.775 |
| **Compartment** | **6032.100** | **3016.040** | **493.703** | **<0.001** |
| Sugar Content | 7.900 | 7.870 | 1.289 | 0.257 |
| **Year** | **164.300** | **164.350** | **26.902** | **<0.001** |
| **Site** | **39.800** | **19.900** | **3.257** | **0.041** |
| **Rootstock x Scion** | **91.500** | **45.750** | **7.489** | **0.001** |
| Compartment x Sugar Content | 4.200 | 2.120 | 0.348 | 0.706 |

Bold values indicate significant results, $P < 0.05$ .

**Table S10.** Linear mixed model results for Chao1 index. Type-3 ANOVA table.

| Factor | Sum Sq | Mean Sq | F value | P |
| --- | --- | --- | --- | --- |
| Rootstock | 2894 | 1447 | 0.815 | 0.445 |
| Scion | 17 | 17 | 0.010 | 0.921 |
| **Compartment** | **772453** | **386226** | **217.635** | **<0.001** |
| Sugar Content | 443 | 443 | 0.250 | 0.617 |
| **Year** | **29154** | **29154** | **16.428** | **<0.001** |
| **Site** | **11230** | **5615** | **3.164** | **0.045** |
| **Rootstock x Scion** | **15806** | **7903** | **4.453** | **0.012** |
| Compartment x Sugar Content | 23 | 12 | 0.007 | 0.994 |

Bold values indicate significant results, $P < 0.05$ .

**Table S11.**
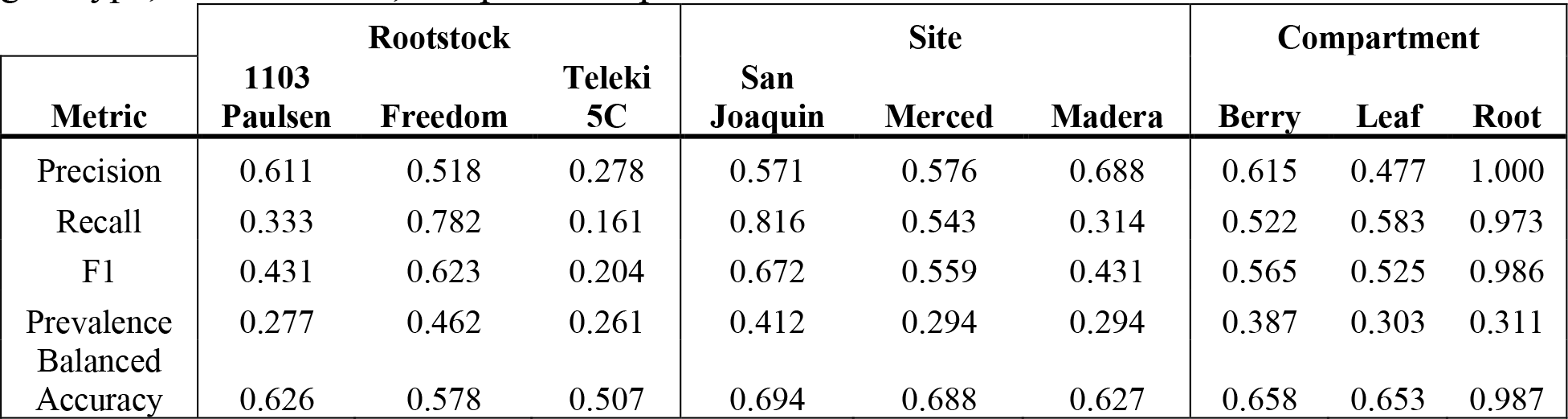
Output statistics for three-class machine learning models predicting rootstock genotype, collection site, and plant compartment.

**Table S12.** Output statistics for binary class machine learning models predicting scion genotype and collection year. In these models Cabernet Sauvignon and 2018 represented the positive class.

| Metric | Scion | Year | Sugar Content |
| --- | --- | --- | --- |
| Precision | 0.704 | 0.824 | 0.742 |
| Recall | 0.559 | 0.949 | 0.857 |
| F1 | 0.623 | 0.882 | 0.795 |
| Prevalence | 0.571 | 0.664 | 0.647 |
| Balanced Accuracy | 0.623 | 0.775 | 0.655 |

